# Computational principles of adaptive multisensory combination in the *Drosophila* larva

**DOI:** 10.1101/2023.05.04.539474

**Authors:** Philip H. Wong, Andreas Braun, Daniel Malagarriga, Jeff Moehlis, Rubén Moreno-Bote, Alexandre Pouget, Matthieu Louis

**Affiliations:** Department of Mechanical Engineering, University of California, Santa Barbara, Santa Barbara, CA 93106, USA; EMBL-CRG Systems Biology Research Unit, Centre for Genomic Regulation (CRG), The Barcelona Institute of Science and Technology, Dr. Aiguader 88, Barcelona 08003, Spain; Center for Brain and Cognition and Department of Information and Communications Technologies, Pompeu Fabra University, Barcelona, Spain; Department of Basic Neurosciences, University of Geneva, Geneva, Switzerland; Department of Molecular, Cellular, and Developmental Biology & Neuroscience Research Institute, University of California, Santa Barbara, Santa Barbara, CA 93106, USA; Department of Physics, University of California, Santa Barbara, Santa Barbara, CA 93106, USA

## Abstract

Many sensory systems have evolved to optimally combine signals from multiple sensory modalities to improve perception. While theories have been proposed to explain how this process is accomplished through probabilistic inference using large neural populations in vertebrates, how animals with dramatically smaller nervous systems such as the *Drosophila melanogaster* larva achieve multisensory combination remains elusive. Here, we systematically characterize larval navigation in different configurations of odor and temperature gradients with optogenetically-controlled noise. Using a data-driven agent-based model, we find that larvae adapt to the reliability of individual sensory signals, and in some cases minimize the variance of the combined signal. Besides firmly establishing that probabilistic inference directs natural orientation behaviors in the *Drosophila* larva, our results indicate that the exact mechanism underlying the combination of sensory information may be modality-dependent. By underscoring that probabilistic inference is inherent to insect nervous systems, our work opens the way for studying its neural implementation.

## Introduction

When confronted with an ever-changing and often perilous environment, how an organism behaves in response to uncertain and incomplete sensory information can be a matter of life and death. Besides the need to assess individual sensory signals accurately, sensory systems must also be able to integrate signals from multiple sensory modalities (e.g. visual, auditory, haptic), some of which may produce conflicting information. This task of “multisensory cue combination” has therefore been the focus of many studies, particularly in psychophysics, to characterize its implementation in different organisms and to evaluate whether these solutions are optimal from a probabilistic point of view (Knill & Pouget, 2004).

One mechanism adopted by organisms to integrate noisy (fluctuating) information arising from different sensory modalities is to prioritize signals based on their relative uncertainty (variance) by using a principle of Bayesian inference. This strategy has the advantages of allowing adaptation to sudden changes in the environment, permitting the filtering of irrelevant information (noise), and improving the signal-to-noise ratio of the combined signal. In humans, for example, the visual-haptic estimation of the height of an object is close to optimal and closely matches the Bayesian estimate (Ernst & Banks, 2002). Similar results have also been observed for other tasks in humans (Hillis et al., 2004), as well as in primates (Gu et al., 2008). To a lesser extent, recent evidence indicates that insect brains may also be capable of implementing similar strategies of cue combination, for example in the integration of directional information in ants (Sun et al., 2020; Wystrach et al., 2015). In addition, the neural integration of multisensory cues has been studied in the adult *Drosophila* and it has been shown that flies are able to dynamically adjust their response to conflicts between visual, olfactory and airflow cues (Currier et al., 2020).

Although the neural implementation of cue combination is not well-understood, various theories speculate about how neural ensembles can implement probabilistic inference (Jordan et al., 2021; Ma et al., 2006). While certain theories require neuronal populations to encode probabilities and information about signal variance (Ma et al., 2006), others suggest the possibility of encoding variability through synaptic plasticity in single neurons (Jordan et al., 2021). Further characterizing multisensory cue combination in a comparatively simple model organism like the *Drosophila* larva is advantageous, not only to reveal how strategies evolve through development, but also to delineate the minimal complexity required to mechanistically implement strategies of multisensory-cue combination (Berck et al., 2016).

While it has yet to be shown how the *Drosophila* larva implements cue combination in natural conditions, previous studies have examined how turns are triggered in the *Drosophila* larva in response to the combination of aversive light input and attractive virtual odor input (Gepner et al., 2015, 2018). In the first study, a computational model that describes the basic transformation of sensory input into turning decisions was built to investigate the sequence of mathematical operations combining multi-modal inputs (Gepner et al., 2015). In subsequent work, a modified version of the same model was used to establish that signals triggering turns adapt to the variance of the individual multi-modal sensory inputs (Gepner et al., 2018). In the present work, we investigate whether this form of variance adaptation fits into traditional cue combination models as observed in other animals and dissect how the mechanism underlying the combination of multi-modal inputs contributes to the overall navigational strategy of the larva. Specifically, we investigate how the *Drosophila* larva responds to gradients of two independent odors, as well as the combination of an odor and a temperature gradient. While chemotaxis and thermotaxis have been studied extensively in the larva (Klein et al., 2015; Louis, 2020; Luo et al., 2010), little is known about how unimodal navigational mechanisms contribute to navigation in unison.

Experimentally, we investigate combinations of thermotactic and olfactory (real and virtual) stimuli in scenarios where cues are directionally similar (congruent) or in opposing (conflicting) directions. Furthermore, we test conditions where noise is added optogenetically to the peripheral olfactory system to study how the combination of multisensory cues adapts to changes in the variance of individual sensory inputs. To capture the precise reorientation mechanisms and navigational behavior of larvae in these scenarios, we built a data-driven agent-based model inspired by Wystrach et al. (2016) that represents both turn rate and turning direction, and models how different sensory inputs are processed and transformed into behavioral outputs. Using this agent-based framework, we tested and simulated different experimental paradigms to narrow down the set of plausible mechanisms for multisensory cue combination in the *Drosophila* larva through a process of elimination. With this approach, we explore computationally how larvae use signal variance to weigh and combine unreliable sensory information from multiple modalities. Using our agent-based model, we conduct a perturbative analysis to characterize the modulatory impact of cue combination on individual aspects of the control of locomotion underlying sensory navigation.

Motivated by a need to go beyond cue-combination models that specifically estimate the properties of a single object (e.g., the width of a bar, (Ernst & Banks, 2002)), we explore different notions of optimality related to sensory navigation in response to realistic configurations of multimodal gradients. Through a generalized formalism of cue-combination strategies, we define a bimodal contrast coefficient that represents the degree to which signal variance is prioritized over the value (reward) of individual signals in the combination of multimodal sensory inputs. In addition to the observation that larvae are near-optimal in both formalisms, we find that their cue-combination strategy can adapt depending on the nature of the sensory information available to the animal.

## Results

### An experimental assay to quantify multisensory combination in the larva

A behavioral assay was developed to study larval navigation in spatial gradients of temperature, a real odor, and a virtual odor induced optogenetically by expressing Chrimson in genetically-targeted olfactory sensory neurons (OSNs). Red light elicited virtual-odor stimulations in the *Or67b*-expressing OSN which is not activated by ethyl butyrate (Kreher et al., 2008; Si et al., 2019)., the real odor used in this study. As a result, the real and virtual odor activated a distinct and independent set of OSNs. In each experiment, larvae at the third developmental instar were uniformly distributed in groups of 10 individuals near the center of a circular behavioral arena coated with agarose (Figure 1A). The motion of the group of larvae was video-monitored during exposure to single or combined sensory gradients. The trajectories of larvae in the arena were then extracted using a custom image processing and tracking software. Larvae were analyzed individually as, given the low density of animals, group effects were found to be negligible in the context of these gradients (see ‘Materials and methods’).

**Figure 1.**
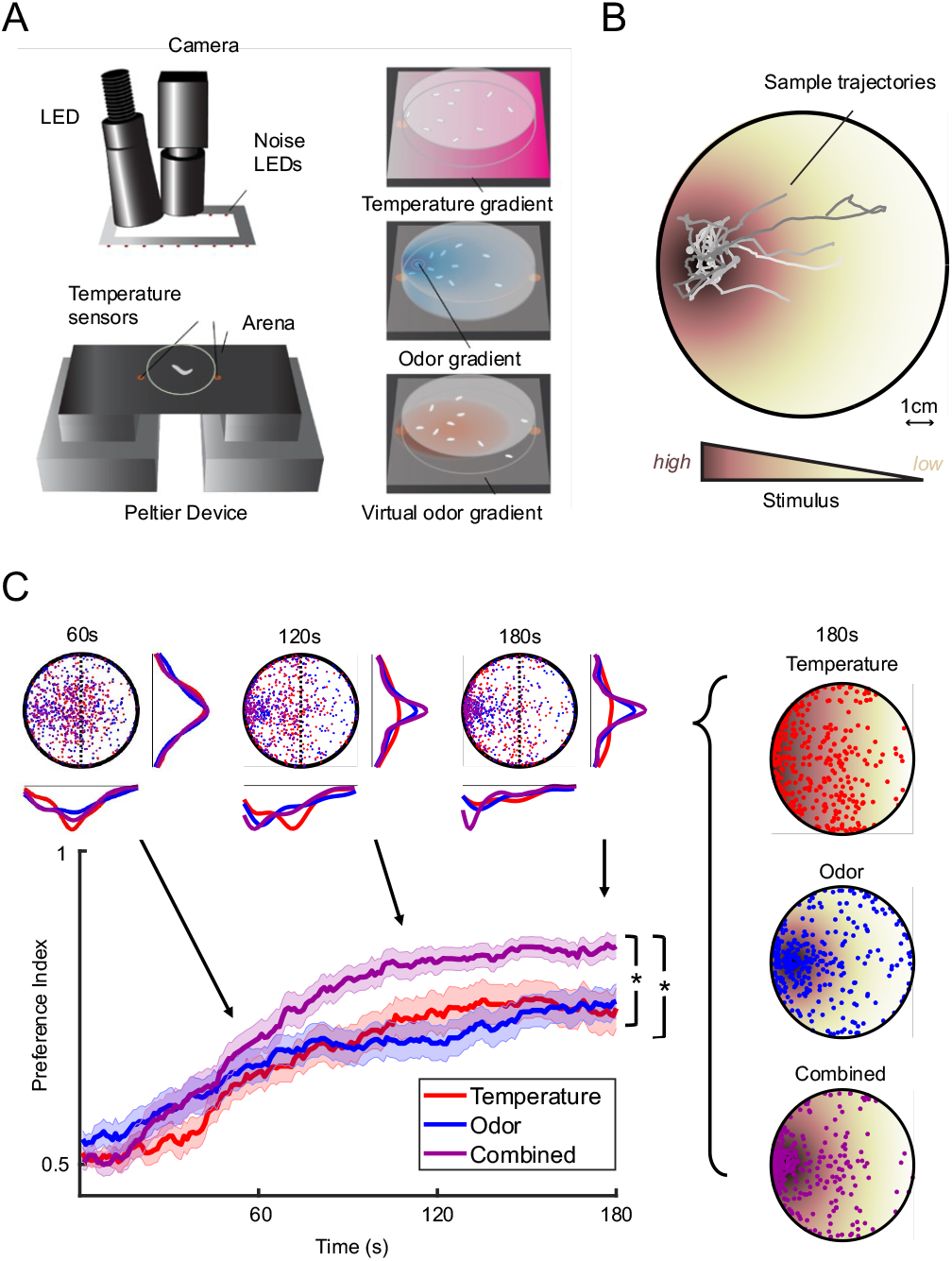
Assay to identify how larvae navigate unimodal (single) and bimodal (combined) gradients. **(A)** Schematic of the behavioral assay, which features gradients of real odor, optogenetically-induced virtual odor, and temperature. **(B)** Representative trajectories of third-instar wild-type (*w*^*1118*^) larvae responding to the combination of an odor and a temperature gradient over a period of 3 minutes. **(C)** Behavioral response of wild-type larvae to the individual odor and temperature gradients and both odor and temperature combined (Odor: Ethyl butyrate, 10^−3^ M; Temperature range: 16-30°C). Larvae were tested in groups of 10 individuals (Odor: n = 27 groups of 10 larvae; Temperature: n = 35; Combined: n = 27). In all subsequent figures, the shaded regions around the preference index curves represent the error bars of the SEM. The asterisks indicate that the preference index of the combined condition was significantly higher than the preference indices of either unimodal condition (after the first minute of the experiment), as assessed using a *t*-test (*p* < 0.025 upon Bonferroni correction). Also illustrated are the overlayed spatial distributions of larvae for each condition at 60, 120, and 180 s (top), and the spatial distributions for each individual condition at 180 s (right).

In conditions where single gradients were presented, which we will refer to as unimodal conditions, larvae navigate unimodal odor, virtual-odor, and temperature gradients by locating the “source”: the region associated with the highest concentration of the attractive odor or the most comfortable temperature in the arena. When placed near the center of the arena, larvae innately navigated to the location of highest odor concentration, highest virtual-odor intensity, or the location with the most preferred temperature, which was slightly higher than 16°C in our experimental conditions (Figure 1B). In the range of temperatures used in the present work, larvae demonstrated robust thermotaxis down temperature gradients toward the coolest region of the arena.

In situations where two gradients are presented at the same time, which we will refer to as bimodal conditions, we initially arranged the gradients in congruent configurations such that both sources were on the same side of the arena with colinear gradients. At the start of the experiment, larvae were placed near the center of the arena and over time distributed in a way similar to the unimodal conditions. Notably, larvae in bimodal conditions demonstrated improved performance in navigating towards the congruent sources compared to the unimodal conditions. For example, the attraction towards the source increased upon combination of an odor and a temperature gradient (Figure 1C). This result is quantified by the preference index, which is the fraction of larvae on the targeted side of the arena (i.e. odor source or preferred temperature) as a function of time:

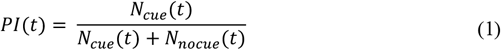

Sluggish larvae displaying an average speed lower than 0.1 mm/s are excluded from the preference index calculation to avoid counting inactive outliers sitting near the starting location. For convenience of notation, we omit the time variable *t*and simply refer to the preference index as the *PI* in the rest of the text. We observed a similar improvement in preference index across all other experimental paradigms with congruent gradients of two distinct odors, a real odor and a virtual odor, as well as a virtual odor and temperature (Figure S1, S2).

### A coarse-grained model suggests that larvae account for cue uncertainty when combining multimodal cues

To characterize how heightened attraction emerges from the combination of olfactory and thermosensory cues in congruent gradients, we started by developing a parameter-free theoretical model using the principle of Bayesian inference to estimate the probability distribution of the positions of individual larvae in the arena (see section Parameter-Free Model in Supplementary methods). The model predicts that the weighting of the information from different gradients is dependent on the uncertainty associated with each gradient. As described in the Supplementary methods, this coarse-grained model estimates the PI of the response to the combined-gradient condition based on the PI of the corresponding unimodal conditions *PI*_1_ *a*nd *PI*_2_:

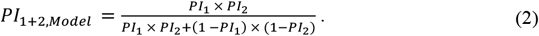

As shown in Figure 2B, we found that the parameter-free model reproduces the behavioral improvement observed in the experimental preference index for the congruent temperature and odor gradient presented in

**Figure 2.**
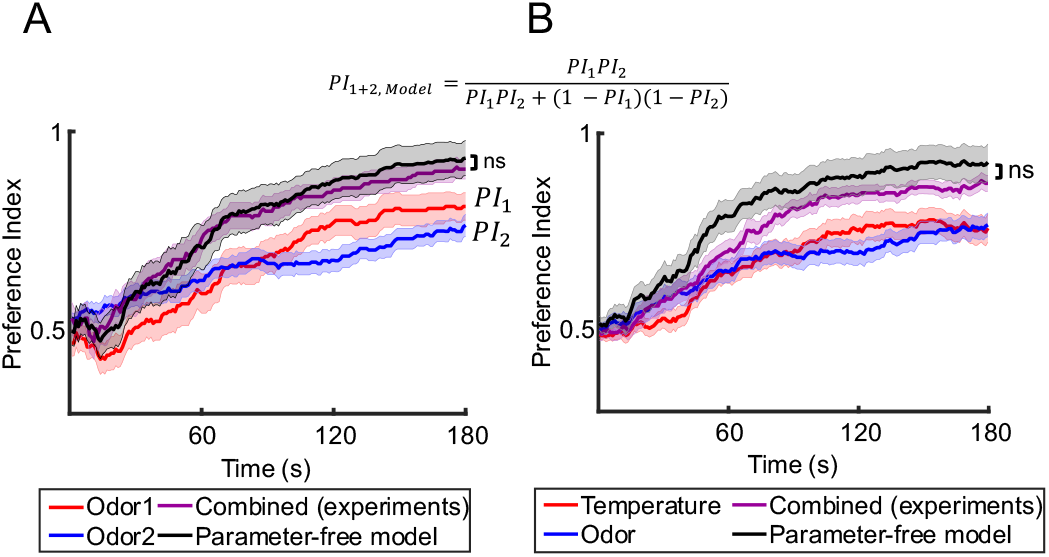
Comparison of experimentally-observed combined preference indices with a coarse-grained parameter-free model for different configurations of congruent gradients. In both configurations, no significant difference exists between the final (180s) preference indices of the experimental data and the parameter-free model (*t*-test, *p* > 0.05). For the full comparison of all congruent gradients tested, see Figure S3. **(A)** Odor + odor (odor 1: 1-hexanol, 10^−2^ M, n = 20; odor 2: Ethyl butyrate, 10^−3^ M, n = 26; Combined: n = 19). **(B)** Temperature + odor, as outlined in Figure 1C.

Figure 1C. In addition, we applied the parameter-free model to predict the behavior of larvae tested in congruent gradients featuring two real odors (Figure 2A), a real and a virtual odor, a real odor and temperature, or a virtual odor and temperature (Figure S3). In all four experimental conditions, the results of the model were in excellent qualitative agreement with the behavior elicited by congruent bimodal gradients, suggesting that real larvae use probabilistic inference to combine sensory information.

### Building an agent-based model to characterize how the combination of sensory cues directs navigation

To analyze the plausibility of different mechanisms of sensory combination and dissect the control of individual reorientation maneuvers, we developed an agent-based model that offers a more realistic description of larval navigation in response to both unimodal and bimodal conditions (Figure 3A). The starting point of our agent-based model is an existing mechanical model of chemotaxis in the *Drosophila* larva (Wystrach et al., 2016), which provides a general framework for describing orientation (“taxis”) behavior elicited by unimodal stimuli. Based on evidence that larvae display continuous lateral oscillations of the anterior body segment during peristalsis, the agent-based model established that a direct sensory modulation of the oscillation amplitude of head-casts could reproduce many signatures of chemotaxis observed in larvae.

**Figure 3.**
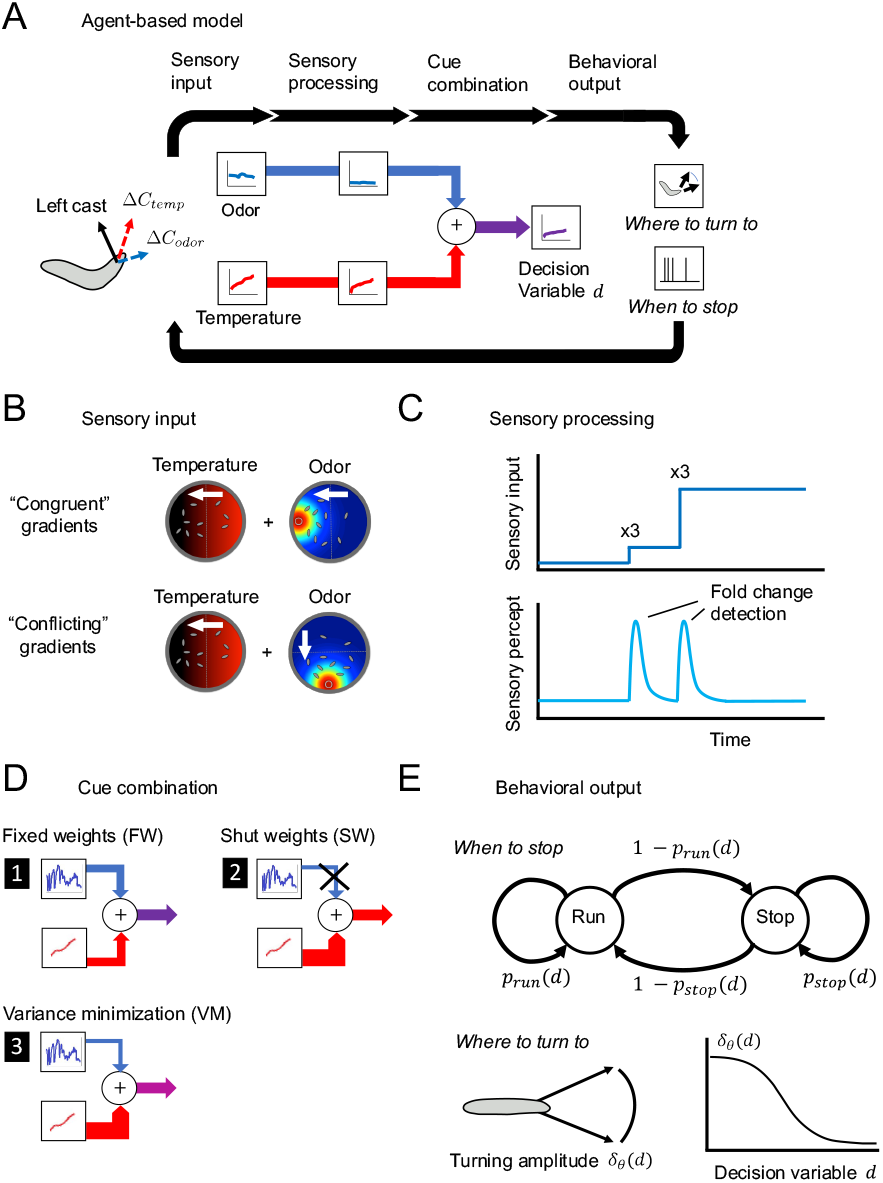
Outline of agent-based model for *Drosophila* larval navigation and set of plausible cue combination models. (**A**) The different stages of the agent-based model are represented in a flowchart from sensory input to behavioral output. The illustration depicts the sensory experience during a left head cast in an odor and a temperature gradien. (**B**) Gradients presented in each experimental paradigm can be congruent (co-linear) or conflicting (90-degree angle). (**C**) Sensory inputs are processed individually with the assumption that the resulting perceptual cue is proportional to relative changes in stimulus strength. (**D**) The perceptual cues from each sensory modality are combined as a weighted linear combination, with weights dependent on the cue combination rule. (**E**) The decision variable determines the amplitude of head casts *δ*_*θ*_ (“where to turn to”) and the probability of mode transitions *p*_*run*_, *p*_*stop*_ (“when to stop”) of the agent larva.

As detailed in the Agent-based Model section of the Supplemental methods, we adapted the model of Wystrach et al. (2016) based on the quantification of our behavioral data to account for a multimodal setting by capturing more closely how different sensory gradients are perceived by the larva, and then by modelling how graded information from two different sensory modalities are combined to drive reorientation maneuvers. In our expanded agent-based model, *Drosophila* larvae alternate between straight runs and directed turns. The alternation between these two behaviors is modulated by the detection of temporal increases or decreases in sensory input. Active sensing is achieved primarily through lateral movements of the head, which assesses the local environment to reorient toward the direction of the gradient. To achieve a realistic representation of the sensorimotor control of larval navigation, we incorporated behavioral mechanisms to describe both how larvae determine *when* to initiate a turn and *where* to turn to.

In the model developed here (Figure S4A), the larva is represented as a single segment from its midpoint to its head — the body segment from the tail to the midpoint is assumed to passively follow the head segment, which is reasonable in first approximation. The agent-based larva may be in one of the following two states: running, where the larva moves at a fixed speed in the direction of its head segment while making small adjustments to its heading, and stopping, where the body segment is stationary but the body segment is free to rotate around the midpoint. The behavioral state of the agent-based larva is updated in discrete time steps. At each time step, the head segment alternates between rotations on the left and the right side of the body axis to mimic the active sampling of sensory conditions surrounding the head. At any given timestep *n*, the larva perceives the sensory input *C*_*n*_ *g*iven by the intensity of the stimulus detected at the tip of the head segment where the olfactory organs are located.

In each experimental paradigm, we simulated the behavior elicited by combinations of real-odor gradients with static virtual-odor gradients or static temperature gradients (Figure 3B). While the profiles of the virtual-odor gradients created with a LED and temperature gradients created with a Peltier element were stationary, the real-odor gradients were created by placing an odor droplet on the side of the source. To simulate the dynamics of the odor gradient during the course of an experiment, we used a biophysical model for the odor diffusion introduced in previous work (Schulze et al., 2015) (see Sensory Stimulus section of the Supplemental methods and Supplementary Video 2).

For each sensory modality presented to the larva, we hypothesized that the resulting percept —the internal representation of the odor— is proportional to relative changes in stimulus strength (Adler & Alon, 2017). More specifically, the model assumes that the perceptual response to the real-odor, virtual-odor, and temperature gradients will be of the form 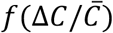, where 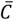 is the background signal level and Δ*C* is the signal difference (Figure 3C). This sensory property is equivalent to Weber law, which has been established in the peripheral olfactory system of the adult fly (Cao et al., 2016; Gorur-Shandilya et al., 2017; Kadakia & Emonet, 2019). We assume that the larval olfactory system detects relative changes in odor concentration, which is supported by the response properties of larval OSNs (Gomez-Marin & Louis, 2012; Schulze et al., 2015) and the apparent concentration-invariance of reorientation maneuvers (Gomez-Marin & Louis, 2012). For temperature, we make the assumption in our agent-based model that the larva perceives relative changes zeroed at the maximum temperature of the behavioral assay (i.e. *C* ← *T*_max_ − *C*). This results in a perceptual response that increases as larvae move away in a temperature gradient from preferred temperatures. Although the sensitivity of the thermosensory system to relative changes has not been explicitly demonstrated, there is evidence that the magnitude of the behavioral response scales with the difference in temperature relative to the deviation from preferred background temperatures (Hernandez-Nunez et al., 2021; Klein et al., 2015). In our simulations, we computed the relative change in stimulus between two consecutive timesteps *n* − 1 and *n* as the following variable:

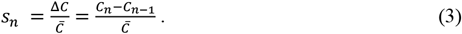

The background signal level 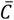 is computed as the midpoint between two timesteps, 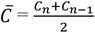. At every time step of the stimulations, the information collected by the two different sensory modalities *s*_1_ and *s*_2_ is combined in a decision variable *d* by using the linear model:

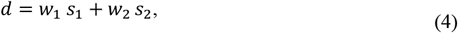

where *w*_1_ and *w*_2_ are weights associated with each cue. Using the model, we examine the three most common weighting strategies, each representing a qualitatively different approach to cue combination:

1. Fixed Weights (FW):

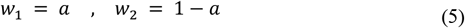
2. Shut Weights (SW):

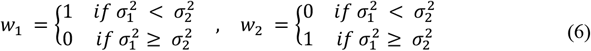
3. Variance Minimization (VM):

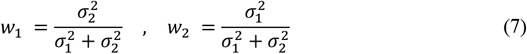

The Fixed-Weights (FW) strategy (Negen et al., 2019) proposes that larvae combine cues with fixed preferences that are independent of the signal variances 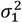 and 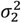. The latter two strategies imply that larvae are also able to adapt their response according to the estimated variance of the sensory inputs accumulated over a time window (for numerical implementation, see Supplementary methods), as established in a previous study (Gepner et al., 2018). Being sensitive to the reliability of sensory inputs is a hallmark of probabilistic inference, a powerful form of computation when dealing with inputs subject to sensory uncertainty. The Shut-Weights (SW) also known as Winner-Take-All strategy (Bresciani et al., 2006; Welch & Warren, 1980) assumes that larvae place absolute priority on the cue that is observed to be more reliable and suppresses the weakest one. The Variance-Minimization (VM) strategy is a linear combination rule that minimizes the variance of the combined signal (Ernst & Banks, 2002). By considering the validity of these three cue-combination strategies for different sensory modalities, we can examine whether variance adaptation is present, and then test the degree to which variance modulates cue combination of multimodal signals.

Finally, the transition rates between the two states, running *p*_*run*_ and stopping *p*_*stop*_, (“*when to stop*”) and the amplitude of head casts *δ*_*θ*_ (“*where to turn to*”) are described as functions of the decision variable *d* using a generalized linear model (Figure 3E, see supplemental methods). The transition probabilities between states and the amplitude of orientation maneuvers are modulated adaptively based on whether the perceived stimulus is attractive (*d* > 0) or aversive (*d* < 0). The direction of head casts alternates at every time step as proposed in Wystrach et al. (2016).

### Application of the agent-based model to explore how sensorimotor integration is implemented in the *Drosophila* larva

The motor parameters of the agent-based model were first optimized to match the behavior of freely foraging larvae. Motor parameters were fit to model the movement patterns of wild-type (*w*^*1118*^) larvae in the absence of any stimulus recorded at high spatio-temporal resolution with the closed-loop tracker from (Schulze et al., 2015) and are assumed to be constants across all experimental conditions. The constants derived from the parameter optimization to model larval motion in the simulations are listed in Table 1 in the Supplementary methods.

The free parameters of the model associated with the multisensory stimuli from each condition (noise, sensitivity) were fit by minimizing the Kullback-Leibler (KL) divergence measured between the spatial distributions and preference indices of simulated and actual larvae. This was achieved by comparing the simulations to actual experimental probability distributions of larvae at different time intervals. We fit the free parameters using the datasets from unimodal conditions (Figure 4A-D). As part of this procedure, the variance associated with each signal was computed using the time course of the stimulus experienced by the agent larva (see Supplementary methods). We then tested each variant of the agent-based model using the three most-common cue-combination rules (Figure 3D) in the combined condition (Figure 4E-F). Based on a process of elimination, we observed that certain cue-combination rules matched the data in some gradient configurations but not others. For example, Figure 4E shows a condition where the experimental PI can be accounted for by the VM rule, but not the FW and SW rules. Additional details about how the models were constrained to capture the behavior of real larvae are provided in Supplementary methods together with Figure S4 and S5.

**Figure 4.**
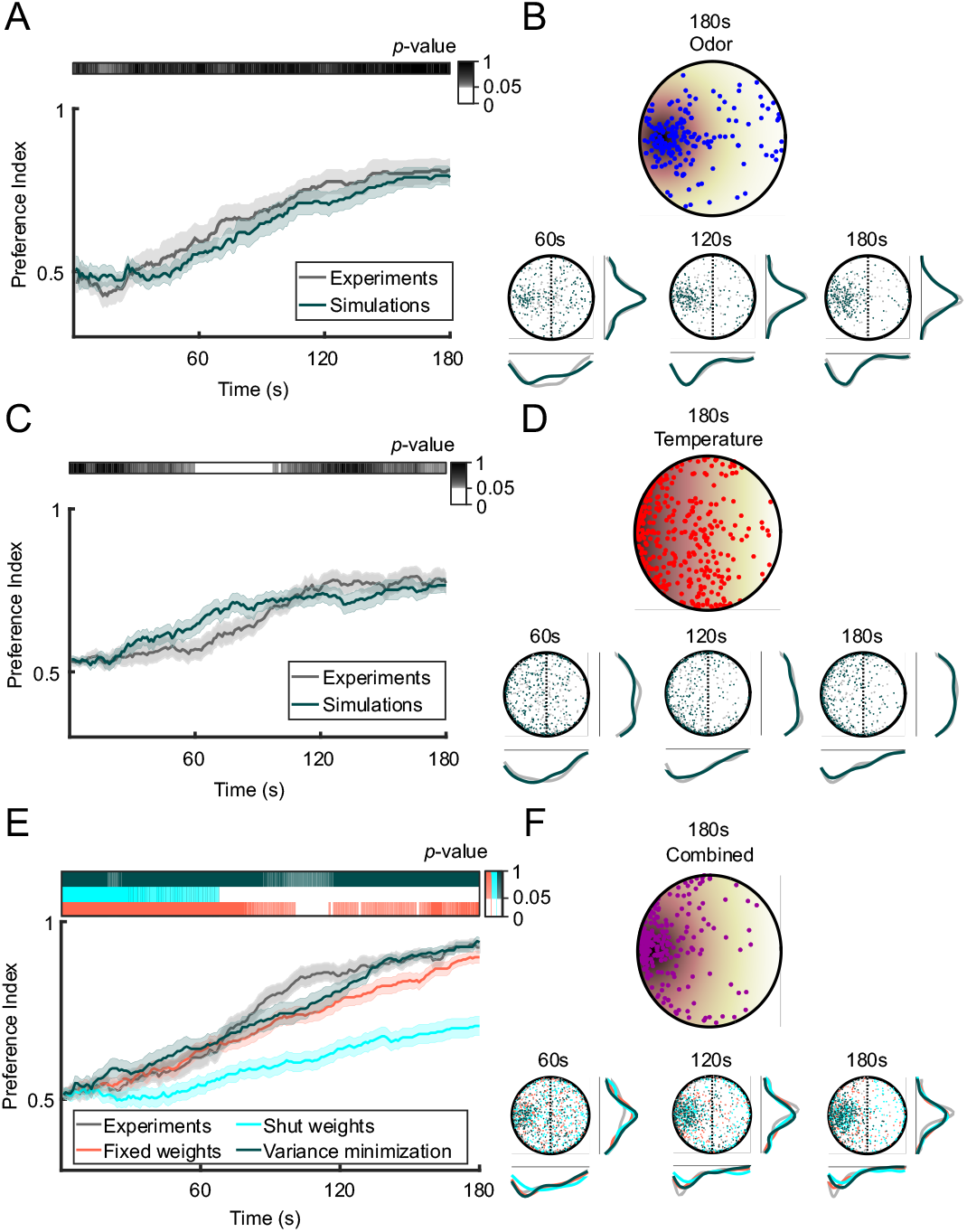
Framework for parameter optimization and testing of the agent-based model for larval navigation. **(A)** Sample simulations for the unimodal odor condition (Odor: Ethyl butyrate, 10^−3^ M) after the free parameters associated with each condition were fit (n = 27). The preference index of simulated larvae was similar to the actual preference indices of wild-type larvae for the entire simulated odor condition (*t*-test, *p* > 0.05). The color bar above the plot indicates the significance of differences between the preference indices of the data and a given fit model. **(B)** The histograms at 60s, 120s and 180s illustrate the spatial distributions of simulated agent larvae and real larvae (gray) for the unimodal odor condition. **(C)** Sample simulations for the unimodal temperature condition (Temperature: 16-30°C) after the free parameters associated with each condition were fit (n = 35). The preference index of simulated larvae was similar to the actual preference indices of wild-type larvae for over 80% of the duration of the simulated temperature condition (*t*-test, *p* > 0.05). The color bar above the plot indicates the significance of differences between the preference indices of the data and a given model. **(D)** The histograms of the spatial distributions of simulated agent larvae (colored) and real larvae (gray) for the unimodal temperature condition. **(E)** Predicted behavioral response of larvae to the combined odor and temperature conditions for each cue-combination rule compared to the actual preference index (n = 27). The preference indices of the simulated Variance-Minimization (VM) and Fixed-Weights (FW) strategies were indistinguishable with the data for over 90% of the entire time course (*t*-test, *p* > 0.05), while the Shut-Weights (SW) strategy remained significantly different from the data after the first minute of the simulation (*t*-test, *p* > 0.05). The color bars above the plot indicate the significant difference between the preference indices of the data and each model. **(F)** Histograms of the spatial distributions of simulated agent larvae (colored) and real larvae (gray) for the combined odor and temperature condition.

We experimentally tested different combinations and configurations of multimodal gradients, including congruent gradients that point in the same direction and conflicting gradients that point in different directions. The KL divergence was used to quantify the degree of similarity between the spatiotemporal distribution of simulated larvae with that of real larvae. By testing paradigms with a variety of gradient geometries, we concluded that the Fixed-Weights model fails to predict behavior in conflicting gradients, such as a conflict between a virtual-odor and a real-odor gradient, (Figure 5A). The Shut-Weights (SW) model underperforms the Variance-Minimization (VM) model in congruent gradients as illustrated with the congruent temperature and real-odor gradient shown in Figure 5B. By comparing the performances on all six experimental paradigms, the VM model gave the most consistent predictions of the three candidate solutions (Figure 5C, bottom panel), even though it did not produce the best fit for all conditions.

**Figure 5.**
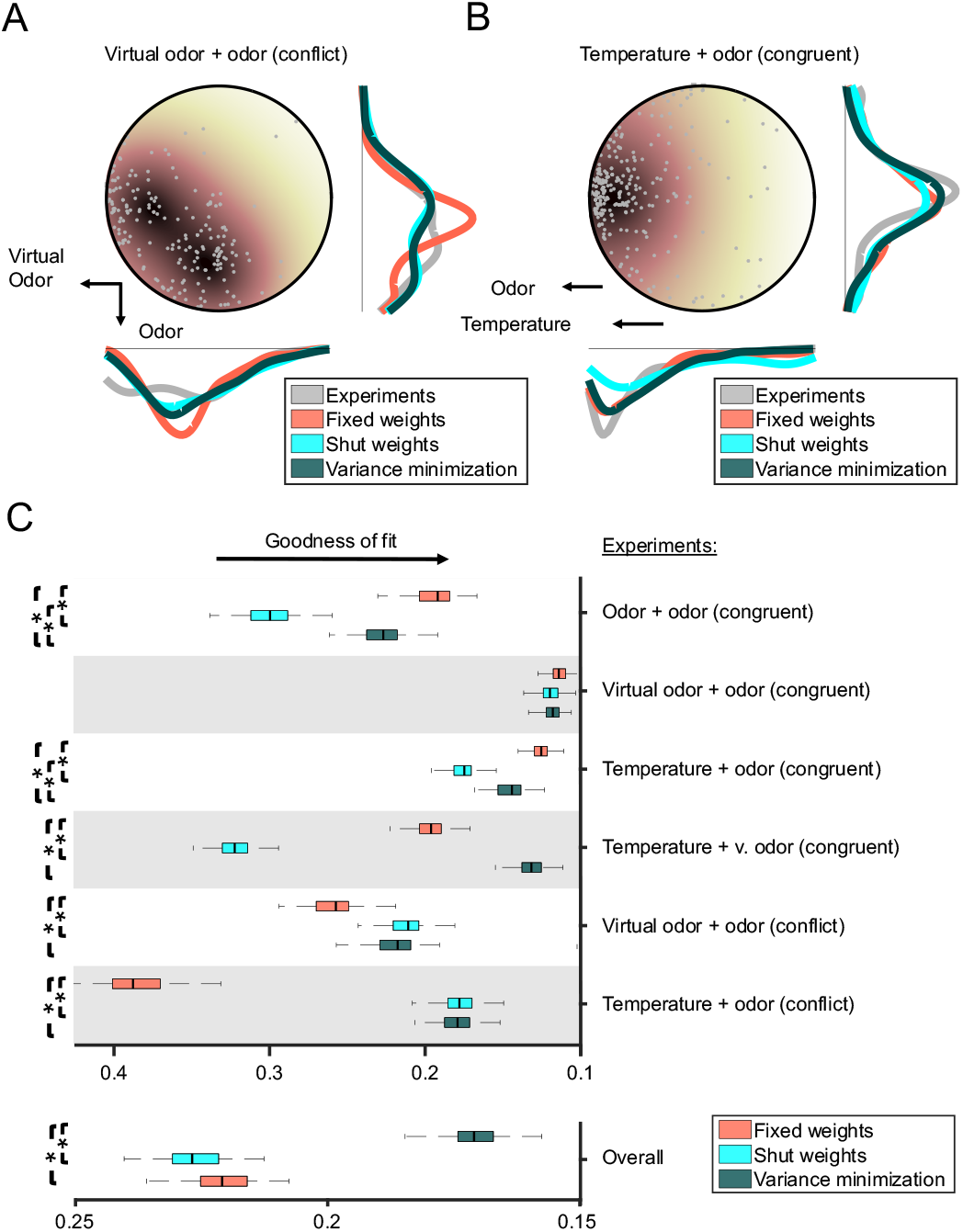
Comparison of the model performances for three cue-combination rules across different experimental paradigms. **(A)** Final distributions of larvae for each simulated cue-combination rule in a conflicting virtual-odor and real-odor gradient (Virtual Odor: *Or67b*>Chrimson, Light 625nm; Real Odor: Ethyl butyrate, 7.5 × 10^−5^M) in comparison to actual *Or67b*-functional larvae (n = 20). The FW strategy led to the poorest fit and was significantly different from both the SW and VM strategies (*t*-test, *p* < 0.05). **(B)** Final distributions of larvae for each cue combination rule in a congruent temperature and odor gradient (Temperature: 20-40°C; Odor: Ethyl butyrate, 10^−3^M) in comparison to actual *Or42a* single functional larvae (n = 30). The SW strategy gave the least accurate predictions and was significantly different from both the FW and VM strategies (*t*-test, *p* < 0.05). **(C)** Comparison of the goodness of fit, as measured by the KL divergence, for cue-combination rules across all experimental paradigms (1-6). The predictions of the VM strategy produced the closest goodness of fit on average to the data (overall), and the VM strategy was significantly different to the FW and SW strategies (*t*-test, *p* < 0.05). Asterisks indicate significant differences between each model to the best fitting model for each experimental paradigm.

Since the VM model combines information with cues that are weighted according to their relative level of reliability (eq. (7)), this scenario suggests that larvae are capable of measuring and processing the variance of their sensory inputs. To test this hypothesis, we experimentally modulated the variability associated with the olfactory cue by optogenetically corrupting sensory encoding in the olfactory sensory neuron (OSN) expressing the *Or42a* odorant receptor, which is tuned to the fruity odor ethyl butyrate (Asahina et al., 2009; Kreher et al., 2008). As described in the Materials and methods, the additive noise consisted in brief random flashes of light inducing the transient depolarization of the *Or42a* OSN expressing Chrimson, while the OSN was responding to the real-odor gradient. As expected, we observed that the chemotaxis of real larvae was weakened when olfactory noise was added to the odor gradient. More surprisingly, we found that thermotaxis improved as quantified by the PI when olfactory noise was added to the detection of a temperature gradient in the absence of any odor gradient (Figure 6A). This seemingly counterintuitive improvement in thermotactic performance illustrates that the weight of each cue is defined by its relative level of reliability: as the noise level increases in the olfactory channel, the reliability of the encoding of genuine dynamic changes due to the odor gradient decreases. In eq. (7), we observe that an increase in *σ*_1_ produces an increase in *w*_2_ irrespective of the presence of any directional signal *s*_1_. Therefore, the injection of pure noise into the olfactory system decreases the weight of this modality and enhances the salience of the thermosensory information.

**Figure 6.**
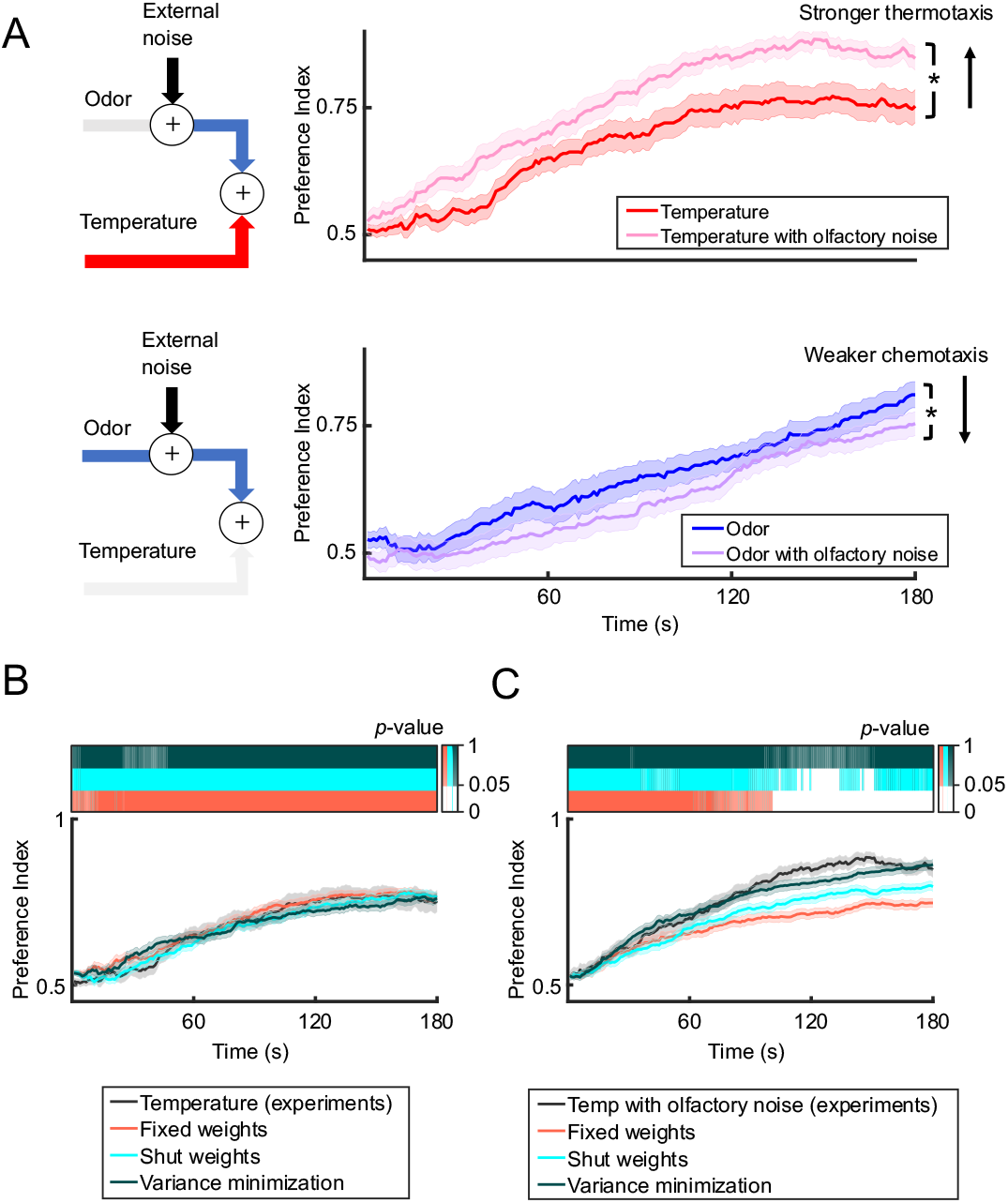
*Drosophila* larvae adapt their orientation responses to the variance of sensory inputs. **(A)** *Or42a*-functional larva navigated odor and temperature gradients while pure noise was injected into the olfactory system via the *Or42a* neuron in the form of optogenetic light flashes. The top graph compares the preference indices for larvae navigating a temperature gradient with and without olfactory noise (Temperature: 20-40°C; Olfactory noise injected through the *Or42a* OSN with light flashes at 625nm, 11.15W/m^2^). The bottom plot compares the preference indices for larvae in an odor gradient versus the same odor gradient with olfactory noise (Odor: Ethyl butyrate, 10^−3^M; Olfactory Noise: *Or42a*, Light 625nm, 11.15W/m^2^). The preference indices for conditions with and without noise are significantly different from one another at the end of the experiment as indicated by the asterisks (*t*-test, *p* < 0.05). **(B)** Actual and simulated response for larvae in a temperature gradient based on the preference index. The FW, SW, and VM strategies are all in agreement with the data for the entire duration of the simulation (*t*-test, *p* > 0.05). **(C)** Actual and simulated response for larvae in a temperature gradient with olfactory noise based on the preference index. The VM strategy is indistinguishable from the data for the entire duration of the simulation (*t*-test, *p* > 0.05), but the FW and SW strategies are significantly different in the latter half of the simulation (*t*-test, *p* < 0.05). The statistical significances of differences between the data and each model are indicated by the color bars above the plots.

To simulate the effects of the olfactory noise on the thermotaxis of agent-based larvae, random disturbances in the activity of the *Or42a* OSN were modeled by the addition of an internal Gaussian noise term to the olfactory signal (see Supplementary methods). In this framework, numerical simulations established that only the VM model was able to qualitatively capture an improvement in thermotactic performances upon injection of pure noise to the olfactory channel (Figure 6B-C). This result strongly supports our hypothesis that the *Drosophila* larva uses an uncertainty-weighted mechanism to integrate multimodal stimuli.

### Two alternative strategies to navigate multimodal gradients optimally

Next, we asked whether the larval nervous system might have evolved to optimize other objectives besides the reliability of each sensory signal to navigate multimodal gradients, and how other strategies might compare to the VM rule (Figure 7). More specifically, we examined whether the exact cue-combination strategy used by larvae is dependent on the nature of the sensory modalities that are combined. Figure 7B illustrates how the VM rule combines a noisy olfactory cue (blue, broader distribution) with mean *s*_2_ and a less noisy temperature cue (red, narrower distribution) with mean *s*_1_ into the decision variable *d*. As a result of eq. (7), the temperature cue has a higher weight than the olfactory cue since *σ*_1_ < *σ*_2_.

**Figure 7.**
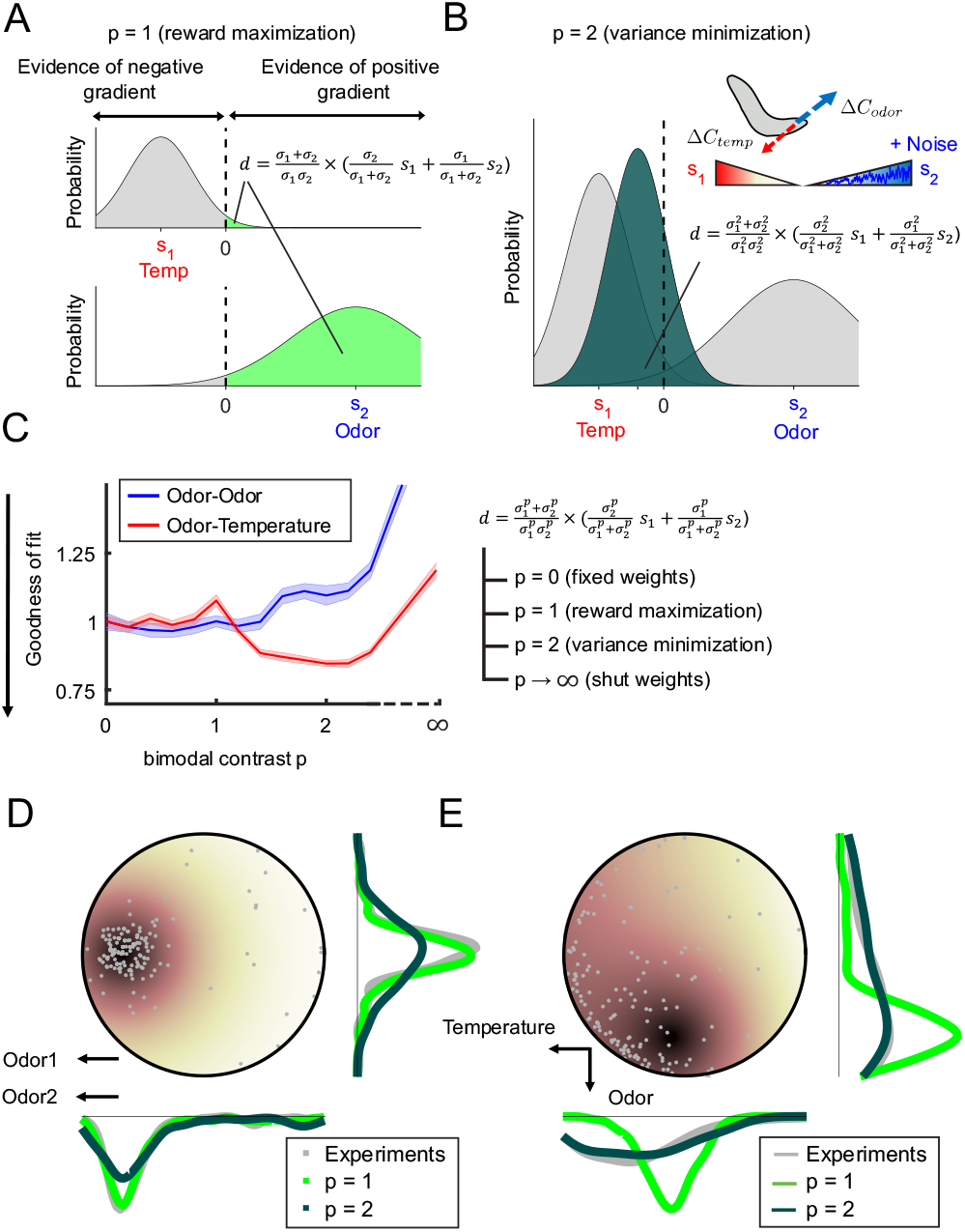
Exploring two different notions of optimality for navigation in sensory gradients. **(A)** Visualization of the reward maximization (RM) rule (p = 1) combining two noisy signals. **(B)** Example of the variance minimization (VM) rule (p = 2) combining a noisy odor signal (blue) and a less noisy temperature signal (red). **(C)** The goodness of fit across experimental paradigms to decision rules with different non-integer values of p. **(D)** Final distributions of larvae in a congruent odor and odor gradient (Odor 1: 1-hexanol, 10^−2^M; Odor 2: Ethyl butyrate, 10^−3^M) for the simulated RM and VM rules in comparison to actual wild-type larvae (n = 19). **(E)** Final distributions of larvae in a conflicting temperature and odor gradient (Temperature: 20-36°C; Odor: Ethyl butyrate, 2.5 × 10^−4^M) for the simulated RM and VM rules in comparison to actual *Or42a*-functional larvae (n = 20).

An alternative objective that a larva could plausibly maximize during navigation is reward. More concretely, we define reward as the probability that motion is directed toward a direction favorable to the encounter of food (motion oriented up an odor gradient) or away from the punishment of potentially noxious heat (motion down a temperature gradient). This strategy, which we call Reward Maximization (RM), is illustrated in Figure 7A with the same two cues configuration presented in Figure 7B. For each of the two cues, the probability that the gradient is positive is equal to the cumulative probability that the cue is greater than zero. Given that the experiments are set up by design for each gradient to be similar in attraction, we make the modeling assumption that there is an equal preference for reaching either favorable sensory condition — whether it is food at the peak of an odor gradient or a temperature range suitable to development. Thus, the reward associated with the maintenance of an ongoing heading is the sum of the probabilities of following a favorable gradient for each of the two modalities. As shown in the Supplementary methods, the sum of these cumulative probabilities can be approximated as the following decision variable:

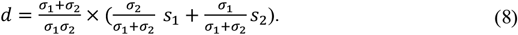

To facilitate a comparison with the reward maximization strategy, the VM rule can be rewritten as:

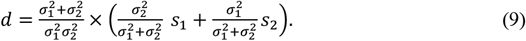

More generally, we note that the VM and RM rules can be written in the form:

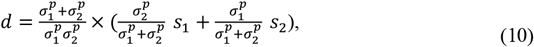

where the value of *p* determines the exact decision rule used. We will hence also refer to the RM strategy as the *p* = 1 rule and the VM strategy as the *p* = 2. Furthermore, the FW strategy can be obtained by setting *p* = 0, while the SW strategy is obtained in the limit as *p* approaches infinity. The decision variable of eq. (10) is generic: it captures a variety of cue-combination strategies defined by the value of a parameter *p* called a *bimodal-contrast parameter*.

### The decision rule applied by a larva is modality-dependent

For a congruent gradient with real odors, the simulated behavior of agent larvae directed by the RM rule reproduced the behavior of real larvae more accurately than agent larvae implementing the VM rule (Figure 7D). This is consistent with the initial results where we showed that the multiplicative combination rule captured the combined PI and that its decision rule corresponds to the case *p* = 1 (eq. (7) in Supplementary methods). On the other hand, the VM rule was more accurate than the RM rule to reproduce larval behavior for a conflicting gradient of odor and temperature (Figure 7E). To generalize this analysis, we set out to compare the goodness of fit of both of the RM and VM rules across all experimental paradigms considered in Figure 5. In addition, we systematically computed the performances associated with specific cases of the decision rule captured by eq. (10), with *p* = 1 representing the RM rule, *p* = 2 representing the VM rule, and the FW and SW rules defining the lower and upper bounds as the value of *p* approaches zero and infinity, respectively. By following this approach, we aimed to determine whether the same rule produced the best fit with the behavior of real larvae for all experimental conditions.

By evaluating the goodness of fit of the simulations to the data for decision rules with different values of *p* (Figure 7C), we made the striking observation that the decision rule applied by real larvae may be dependent on the sensory modalities being combined. While experimental paradigms combining odor and temperature gradients were on average best predicted by decision rules with a value of the bimodal-contrast parameter *p* close to 2, experimental paradigms combining two odor gradients had a goodness of fit curve that suggested the use of a decision rules with a bimodal-contrast parameter close to 1.

To understand why *Drosophila* larvae may use different cue combination strategies depending on the environmental context, we turned to numerical simulations. We quantified how well agent larvae navigated toward favorable gradients using each strategy. To compare how the *p* = 1 rule (equivalent to RM) performed with respect to the *p* = 2 rule (equivalent to VM), we defined two additional metrics quantifying larval behavior to explore and reveal the nuances between the two strategies (Figure 8A-B). The first is “*Reward*”, which would presumably be maximized under the *p* = 1 rule; the second is “*Fraction at Source*”, which is a generalization of the PI beyond congruent gradients. The “*Fraction at Source*” metric, like the PI, quantifies the proportion of larvae that are within specified regions defining favorable conditions (peak of the odor gradient or region with a comfortable temperature, see Supplementary methods). The “*Fraction at Source*” metric is binary: either an animal is inside or outside a favorable region. The “*Reward*” metric defines in a graded way how well larvae remain near or at a favorable location on average. For conflicting gradients, the *Reward* metric can take relatively large values when a larva is located in a region representing a trade-off between the odor and the temperature gradients, whereas the *Fraction at Source* metric leads to 0 values unless the larva has focused on one of the two gradients. Thus, these two metrics tell us how effective each cue combination strategy is at achieving a trade-off between two gradients.

**Figure 8.**
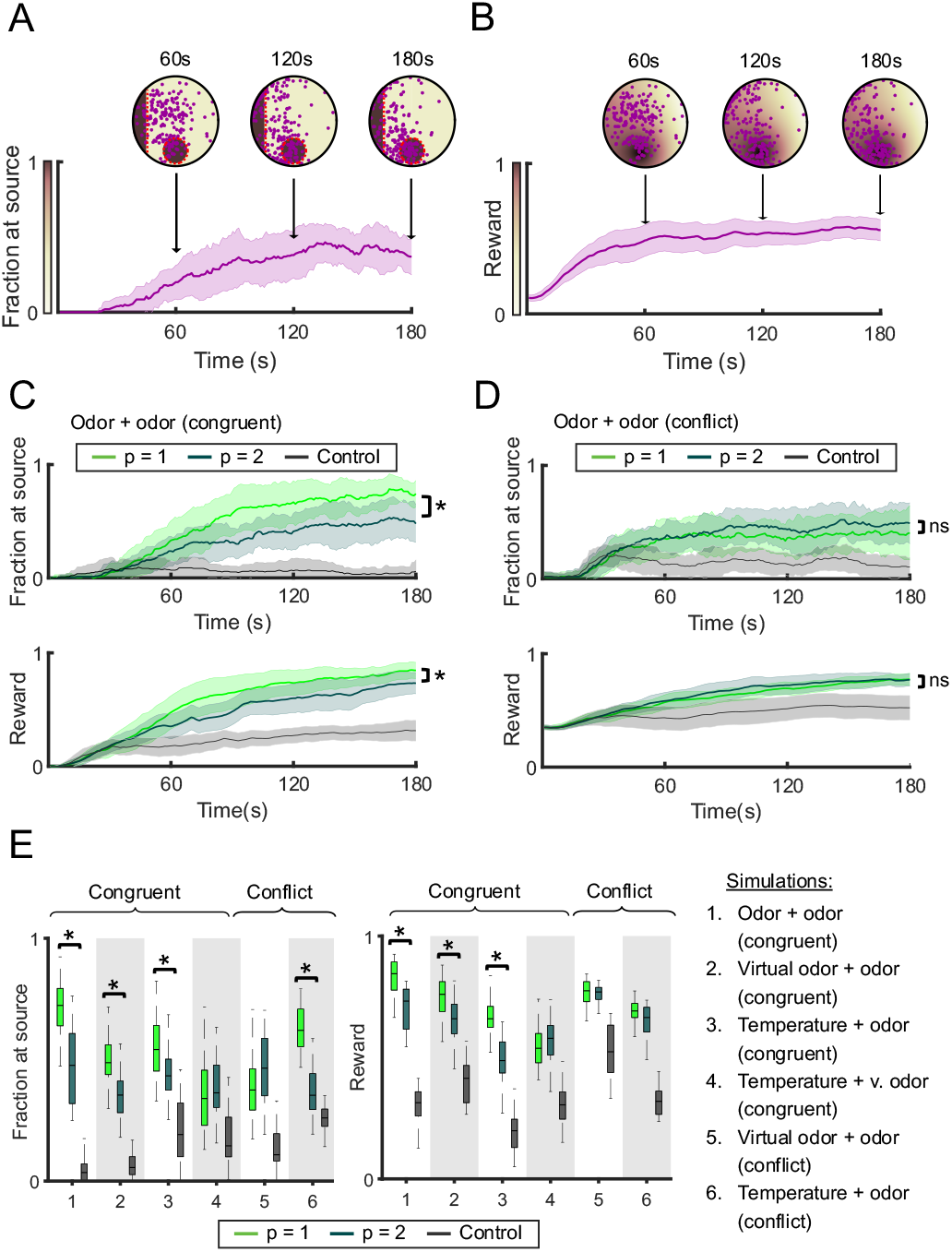
Comparison of the overall performances and characteristics of the RM rule (*p* = 1) and the VM rule (*p* = 2) directing the behavior of simulated agent-based larvae. **(A)** Metric quantifying the **“***Fraction at Source*” metric to quantify how well larvae remain near the source for conflicting temperature + odor gradients. Red dotted lines indicate the boundaries of the two sources. The color gradient indicates the performance of larvae at each location in the arena. **(B)** Metric quantifying the “*Reward*” for the same data as panel A. In both panel A and B, a higher score implies a better performance. **(C)** Comparison of the *Fraction at Source* and *Reward* for a pair of congruent odor + odor gradients. The control condition refers to the performance of simulated agent-based larvae in the absence of any sensory (C-E) information (i.e., decision variable *d* = 0). Simulations of the RM and VM rules lead to a significant difference in both the final *Fraction at Source* and *Reward* (*t*-test, *p* < 0.05). **(D)** Comparison of the *Fraction at Source* and *Reward* metrics for a pair of conflicting odor + odor gradients. Both rules result in a significant difference in the final *Fraction at Source* (*t*-test, *p* < 0.05) but not the reward (*t*-test, *p* > 0.05). **(E)** Comparison of the *Fraction at Source* and *Reward* across all experimental paradigms. The RM rules and VM rules were significantly different for all conditions by both metrics (*t*-test, *p* < 0.05/6 upon Bonferroni correction) except for conditions with conflicting gradients. The asterisks indicate significant differences between the RM rule (p = 1) and the VM rule (p = 2) for each condition.

When we applied the two metrics to quantify the behavior of simulated agent larvae directed by the *p* = 1 (RM) and *p* = 2 (VM) rules, we observed that the differences between the two rules were more significant in congruent gradients than in conflicting gradients (Figure 8C-D). The reward gained by using *p* = 1 instead of *p* = 2 was more significant for congruent gradients compared to conflicting gradients (Figure 8E). We also numerically validated this effect through simulations of a fictive scenario where the conflict angle was sequentially modulated from 0 to 90 degrees (Figure S8). This hints that the advantages of *p* = 1 over *p* = 2 are situational. When comparing these metrics across experimental paradigms, we observed that in general, the *p* = 1 rule performs equally well or better than *p* = 2 when it comes to maximizing the net reward that arises from the combination of two modalities. Effectively, the RM rule achieves a tradeoff between the hedonic value associated with each sensory gradient.

## Discussion

In the present work, we developed an experimental paradigm to quantify the behavior of larvae experiencing congruent or conflicting spatial gradients of odor and temperature. Using this paradigm, we demonstrated that larvae are capable of adjusting the sensitivity of individual sensory channels to changes in the variance of signals transmitted by each modality. In a similar vein as the model delineated in (Gepner et al., 2018) for larvae stimulated by nondirectional white noise with different statistical properties, we establish that the mechanism for variance adaptation can also be described as a weighted sum of sensory cues with weights modulated by signal variance.

While previous work in the larva analyzed multisensory combination mechanisms by observing one specific behavior — the “*when to turn*” mechanism that controls the timing of sensory-driven transitions from running (crawling) to turning (Gepner et al., 2015, 2018), we extended this analysis to directional cues and showed that variance adaptation generalizes to the navigation algorithm as a whole including the mechanism of “*where to turn to*” that creates a turning bias towards favorable sensory gradients. Through numerical simulations, we used a data-driven agent-based model to establish that both of these orientation mechanisms are necessary to account for the navigation of real larvae in multimodal stimuli as removing either component leads to a reduction in performance (Figure S5E). Similar to the adult fly (Demir et al., 2020), the ability to bias turning toward the gradient (“*where to turn to*”) was found to be critical for larvae to navigate toward and accumulate near the odor source.

We tested different plausible strategies for combining sensory inputs, starting with a comparison between the Variance-Minimization (VM), the Fixed-Weights (FW) and the Shut-Weights (SW) rules. The FW and SW rules can be viewed as opposite extremes in the framework of Bayesian cue integration (Ernst & Banks, 2002): while the FW rule always integrates both sensory stimuli, the SW rule systematically discards the less reliable sensory stimulus. This explains why the SW rule is sometimes called Winner-Take-All rule. Similar comparative approaches have been used in the past to compare and evaluate how well different cue combination models fit behavior, for example in human behavior in a two-alternative-forced-choice task (Weisswange et al., 2011). In our results across experimental paradigms, the VM rule accounted best for the behavioral data, while we found that the FW and SW rules were insufficient on their own to adequately reproduce the navigational behavior of larvae for all tested conditions. Next, we introduced the Reward-Maximization (RM) rule, which differs from the VM rule in that it does not assume that the two gradients originate from the same object and location, and seeks to maximize the expected reward of the two gradients (see Figure 7A-B and Supplementary methods). Given the assumptions of the model, both the VM and RW rules are optimal with respect to the objectives they seek to maximize: in the case of the RM rule, it is the reward —strength of the odor stimulus and comfort level of the temperature— that is optimized whereas in the case of the VM rule, it is the reliability of the combined signal.

Since the cue-combination strategies compared in the present study could simply represent four mechanisms out of a limitless set of possible models, we developed a framework to map all four models into a canonical model described in eq. (10) defined by the value of a *bimodal-contrast parameter p*. With this generalized set of models, we showed that our results remained the same in that the RM (*p* = 1) and VM (*p* = 2) were most representative of the way cue combination is implemented by real larvae. Furthermore, we found that some experimental paradigms were better accounted for by the RM rule while others appeared to be more compatible with the VM rule, depending on the pairs of sensory modalities combined by the animal. In particular, the behavior of larvae in a real-odor gradient combined with a congruent temperature gradient was better explained by a principle of variance minimization (VM rule). We believe that this gradation in the decision rule across sensory modalities might reflect the existence of different noise-suppression mechanisms on the underlying behaviors.

Intuitively, larvae may have developed mechanisms of sensory cue combination resembling the RM and the VM rules to exploit different aspects of the sensory conditions that favor their survival in complex natural environments. This hypothesis was tested numerically by evaluating the performance of simulated agent larvae directed by either of the RM (*p* = 1) and VM (*p* = 2) rules in each experimental paradigm (Figure 8C-E), as well as in hypothetical scenarios not tested with real larvae (Figure S8) that include more realistic three dimensional environments. Not surprisingly, we found that larvae experienced a larger “reward” on average with the RM (*p* = 1) rule compared to the VM (*p* = 2) rule. However, the comparison between the RM and VM rules led to more ambiguous results when performances were evaluated based on the fraction of larvae reaching the “source”, as differences in performances between the two rules vanished in conflicting gradients compared to congruent gradients. This result is consistent with the fact that increasing the spatial proximity between cues leads to a smaller improvement in signal reliability during cue combination (Gepshtein et al., 2005).

In the extreme scenario where gradients are pointing at a 90-degree angle, both the RM and VM rules perform similarly as the combination of sensory information becomes less advantageous (Figure 8D and Figure S8A-B). In addition, the two rules differ in that the RM (*p* = 1) rule is closer to the FW (*p* = 0) rule, which always integrates information from both sensory inputs. By contrast, the VM rule leads to a choice of one source over the other resembling the SW (*p* = ∞) rule. When presented with two sources of sensory information, virtual larvae using the RM rule were more prone to remain in between two attractive sources while larvae using the VM rule tended to choose one source over the other. Our agent-based model provides a computational platform to investigate larval integration strategies in more realistic settings, such as navigation on the surface of a sphere (i.e. a rotting piece of fruit). For example, we find that our results extend to a conflict between two attractive odor sources on a spherical surface (Figure S8C).

To explain why larvae appear to utilize the more-integrative RM (*p* = 1) rule in odor-odor gradients but use the choice-like VM (*p* = 2) rule in odor-temperature gradients, we speculate that this nuance may be an example of bet hedging, when organisms suffer decreased fitness in comfortable conditions in exchange for increased fitness in stressful conditions (Danforth, 1999). A larva that cannot feed in a region of moderate temperature is less likely to survive than a larva that chooses to either follow an odor gradient predictive of the presence of food even at the cost potential of noxious heat or to navigate toward a cooler region where food might be found eventually. In the case of odor-odor gradients, larvae might have an advantage to combine multiple chemical cues in a more integrative way given that food sources typically release dozens or hundreds of distinct odorant molecules that are detected by the peripheral olfactory system. By contrast, in situations that present possible danger like aversively high temperatures or starvation in the absence of food, it may be more prudent for larvae to select the more reliable sensory modality earlier as predicted by the VM rule.

Here, we report experimental and modeling-based evidence that *Drosophila* larvae are capable of computing and combining the reliability of sensory inputs to organize orientation behavior in natural conditions. This result suggests that the nervous system of organisms as simple as the *Drosophila* larva can achieve probabilistic inference —a form of computation highly advantageous in uncertain environments. Moreover, the ability of the larva to adapt its navigation strategy to the nature of the perceived multisensory signals offers an opportunity to study differences in the neural implementation of two general rules achieving cue combination based on probabilistic inference, reward maximation and variance minimization. With the availability of the larval brain connectome (Winding et al., 2023), the *Drosophila* larva sets a path to pinpoint where and how different sensory cues are combined and to investigate how these rules evolve across different development stages, such as for the cue integration of odor and wind in the adult fly (Currier et al., 2020; Matheson et al., 2022).

## Acknowledgements

We thank Primoz Ravbar, Jane Loveless, and Ann Hermundstad for their comments on the agent-based model. We thank the Jan Drugowitsch and the Mainen lab for discussion during an early phase of the work. This work was funded by a HFSP Research Grant (RMB, AP and ML), a La Caixa fellowship (AB) and startup funds from the University of California, Santa Barbara (PW and ML).

## Material and Methods

### Fly Stocks

Fly stocks were raised in a 12h light-dark cycle at 22°C/60% humidity. All behavioral experiments were conducted with third-instar larvae reared for 120 hours in tubes on conventional cornmeal-agar fly food. Before each experimental test, larvae were separated from the food by rinsing with a 15% (wt/V) sucrose solution according to a previously established protocol (Louis et al., 2008; Schulze et al., 2015). Testing occurred between 30 to 120 minutes after the introduction of the sucrose. The *w*^*1118*^ strain was used as “wild-type” larvae in experiments combining real odor and temperature gradients. For experimental paradigms involving optogenetically induced virtual odor gradients, the *w*;+;*Or67b*-Gal4 and *w*;*Or42a*-Gal4;+ strains were used to drive the expression of Chrimson in single OSNs. Odor-virtual odor experiments were performed with *w*;+;*Or67b*-Gal4 larvae, while temperature-virtual odor experiments were achieved with *w*;*Or42a*-Gal4;+ larvae.

### Behavioral Assay

The behavioral assay was built using two Peltier elements (CPP-065, TE Technology Inc., USA) attached to a rectangular copper plate via thermo-conductive paste (Céramique, Arctic Silver, USA). Between the Peltier elements, two temperature sensors (Thermistor: MP-2444, TE Technology Inc., USA) were embedded into the metal plate. The temperature of every sensor was monitored by a separate control unit that regulated the Peltier element. Linear temperature gradients were established by setting different target temperatures at each sensor. For an independent temperature assessment, a thermometer with a surface probe (MM2000, TME Electronics, UK, and TS01-S, Surface/Immersion Probe Backfilled, TME Electronics, UK) and an infrared thermometer (Fluke 561, Fluke, USA) were also used to confirm the linear temperature gradient experienced by animals on the surface of the behavioral arena. Virtual odor gradients and noise were generated using red light (62 5nm) by LEDs mounted above the assay (PLS-0625-030-S, Mightex Systems, Canada). The emitted light passed through a mask (exponential cone r=16.5 mm diameter, Leicrom, Spain) in front of the LED resulting in a light gradient in the behavioral arena. In noise experiments, light flashes illuminating the behavioral arena evenly were added on top of the presented gradients. These flashes originated from a rectangular array of red LEDs (Flexible LED strip red 30 x SMD-LED, 850 nm, 12 V, Lumitronix, Germany). Odor gradients were established by pipetting 5 μL of an odorant dilution into a transparent reinforcement ring at the bottom of the arena. In each experiment, the circular behavioral arena was coated with a slab of 3% agarose with a diameter of 107mm. A camera (Stingray F145B ASG, Allied Vision Technologies GmbH, Germany) recorded the behavior of the group of ten larvae for 300 seconds at seven frames per second. An infrared filter (Optical Cast Plastic IR Longpass Filter, Edmund Optics, USA) was placed in front of the camera to exclude any light artifacts.

### Tracking of Animal Posture and Behavioral Quantification

Larvae were tracked offline with a custom-written software in MATLAB. Individual video frames were processed using a black-and-white threshold to perform background subtraction and a size threshold to identify larvae-sized objects. The identities of larvae were labelled in the first frame of the experiment, and subsequent labels were assigned both automatically and manually. The distances between tagged larvae in neighboring frames were computed to match larvae from one frame to the next.

### Parameter optimization and performance quantification of the agent-based model for larval navigation

The constants defining larval navigation in the absence of sensory stimuli (i.e. *d* = 0) were fit using maximum likelihood estimation (Figure S5A-5D). The resulting running, stopping, and head-casting statistics generated by our model were in agreement with actual unstimulated larvae from the closed loop tracker built in (Schulze et al., 2015). To define an appropriate level of complexity for the model, the Akaike information criterion (AIC) and Bayesian information criterion (BIC) were used to quantify the relative importance of each variable in describing larvae behavior in each experimental paradigm. This approach was used to select a final agent-based model with enough degrees of freedom to recapitulate larval navigation across different gradient configurations (Figure S5F). These parameters were then tuned to each experimental paradigm using the unimodal conditions as training data. We defined the objective function to be minimized as the mean Kullback-Leibler divergence (Kullback, 1951) between the simulated and actual X, Y spatial distributions of larvae over the entire time course of the experiment. The parameter sets for each experimental paradigm were optimized using the Global Optimization Toolbox in MATLAB.

To compare larval performance between the Reward Maximization (*p* = 1) and Variance Minimization (*p* = 2) rules, we defined two metrics: “*Fraction at Source*” and “reward”. The *Fraction at Source*, like the preference index, computes the fraction at larvae within bounded regions near the peak of each gradient that is present. Reward assigns each larva a score based on its sensory experience relative to the peaks of each gradient that is present, situated between 0 – the worst location possible and 1 – the best location (see Supplementary methods).

## Supplementary Information

**Figure S1.**
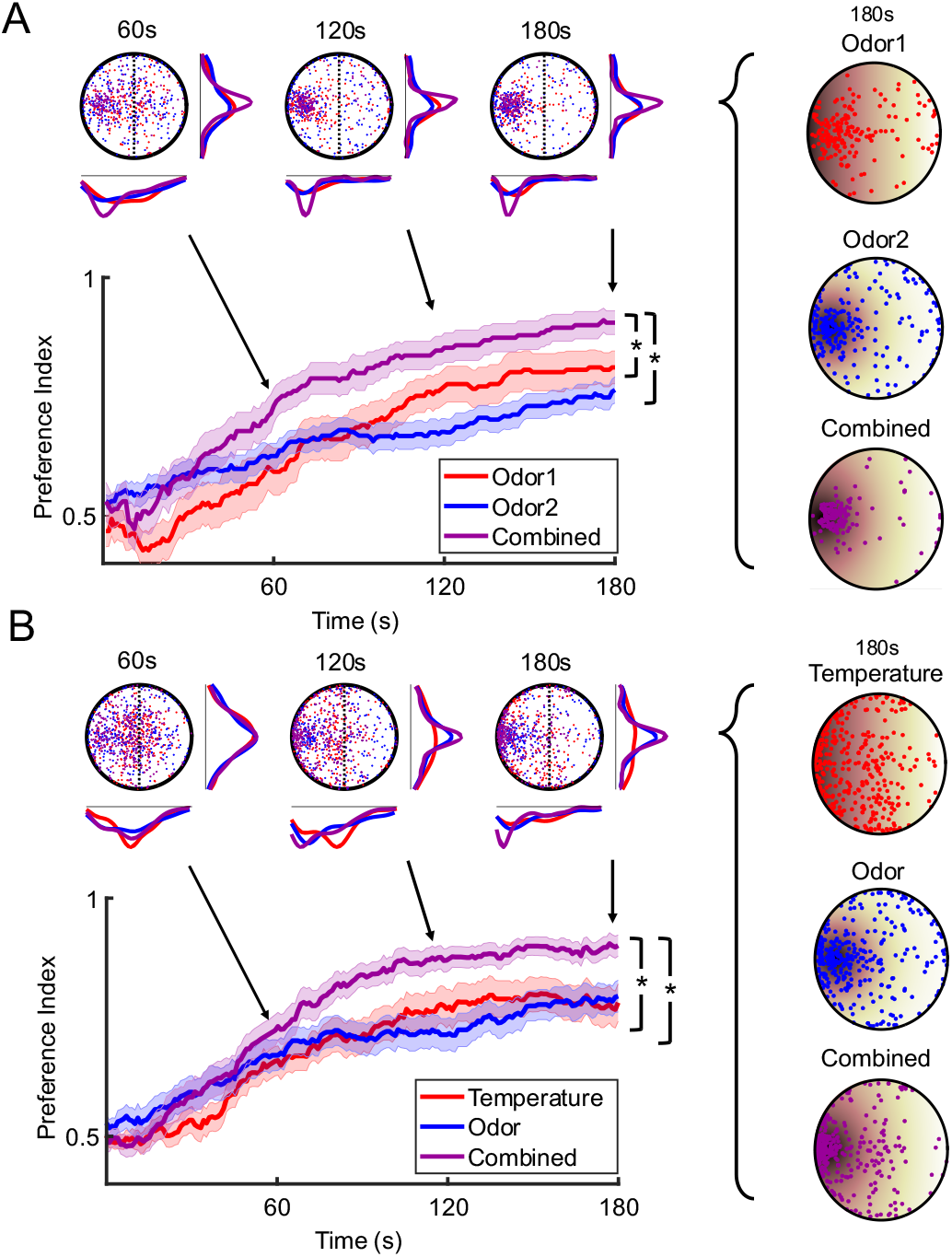
Preference indices corresponding to the performances of wild-type larvae for congruent gradients: odor + odor and odor + temperature. When two congruent unimodal gradients are combined, the final preference index is significantly higher than the preference indices of either unimodal condition as indicated by the asterisks (*t*-test with Bonferroni correction, *p* < 0.025). The shaded regions around the preference index curves indicate the error bars of the SEM. **(A)** Odor + odor (odor 1: 1-hexanol, 10^−2^ M, n = 20 groups of 10 larvae; odor 2: ethyl butyrate, 10^−3^ M, n = 26; combined: n = 19). **(B)** Temperature + odor (odor: ethyl butyrate, 10^−3^ M, n = 27; temperature: 16-30°C, n = 35; combined: n = 27).

**Figure S2.**
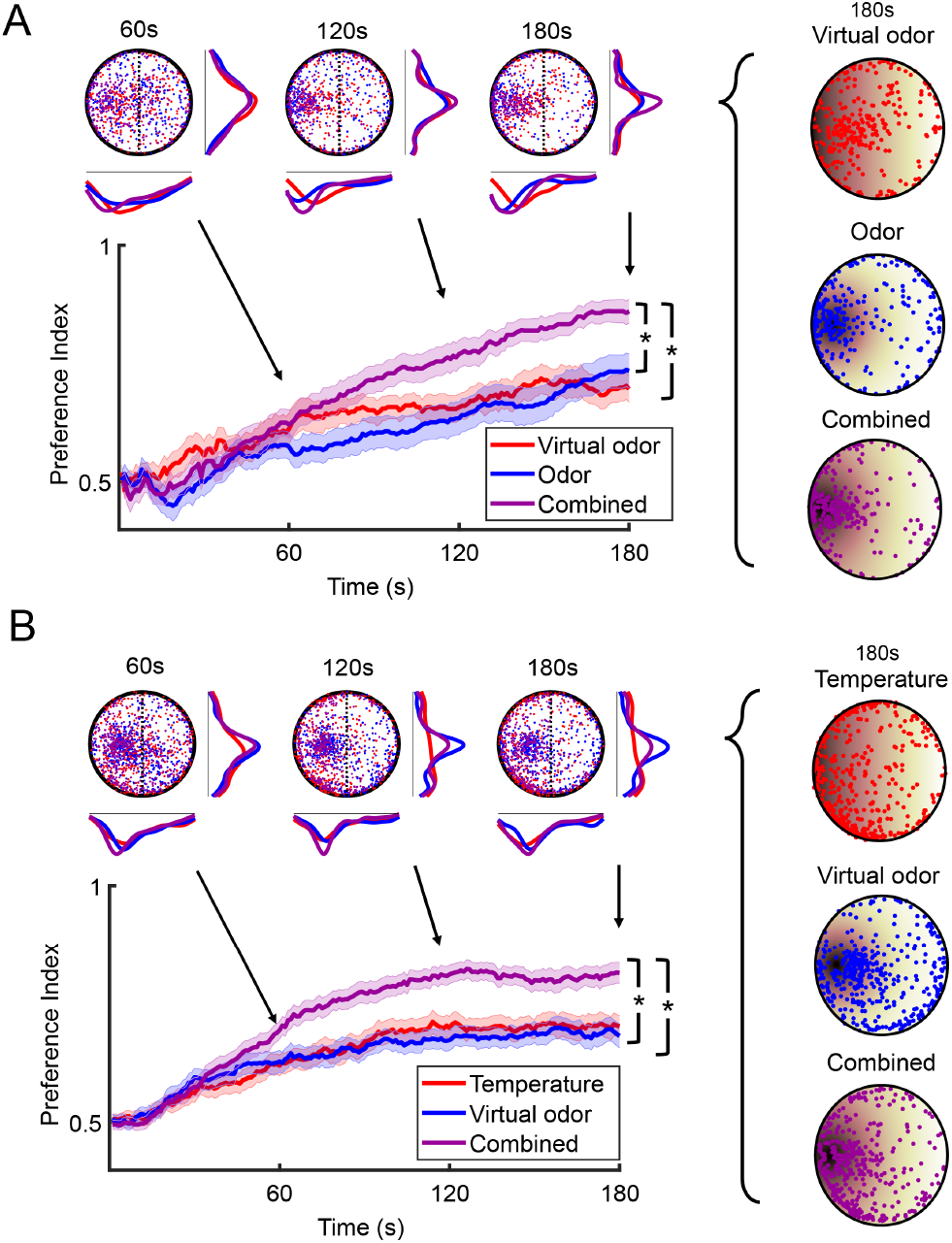
Preference indices corresponding to the performances of wild-type larvae for congruent gradients: virtual odor + odor and virtual odor + temperature. When two congruent unimodal gradients are combined, the final preference index is significantly higher than the preference indices of either unimodal condition as indicated by the asterisks (*t*-test with Bonferroni correction, *p* < 0.025). **(A)** Virtual odor + odor (virtual odor: *Or67b*>Chrimson, light 625nm, n = 30; real odor: ethyl butyrate, 2.5 × 10^−4^ M, n = 30; combined: n = 30). **(B)** Temperature + virtual odor (virtual odor: *Or42a*>Chrimson, n = 49; temperature: 20-40°C, n = 49; combined: n = 49).

**Figure S3.**
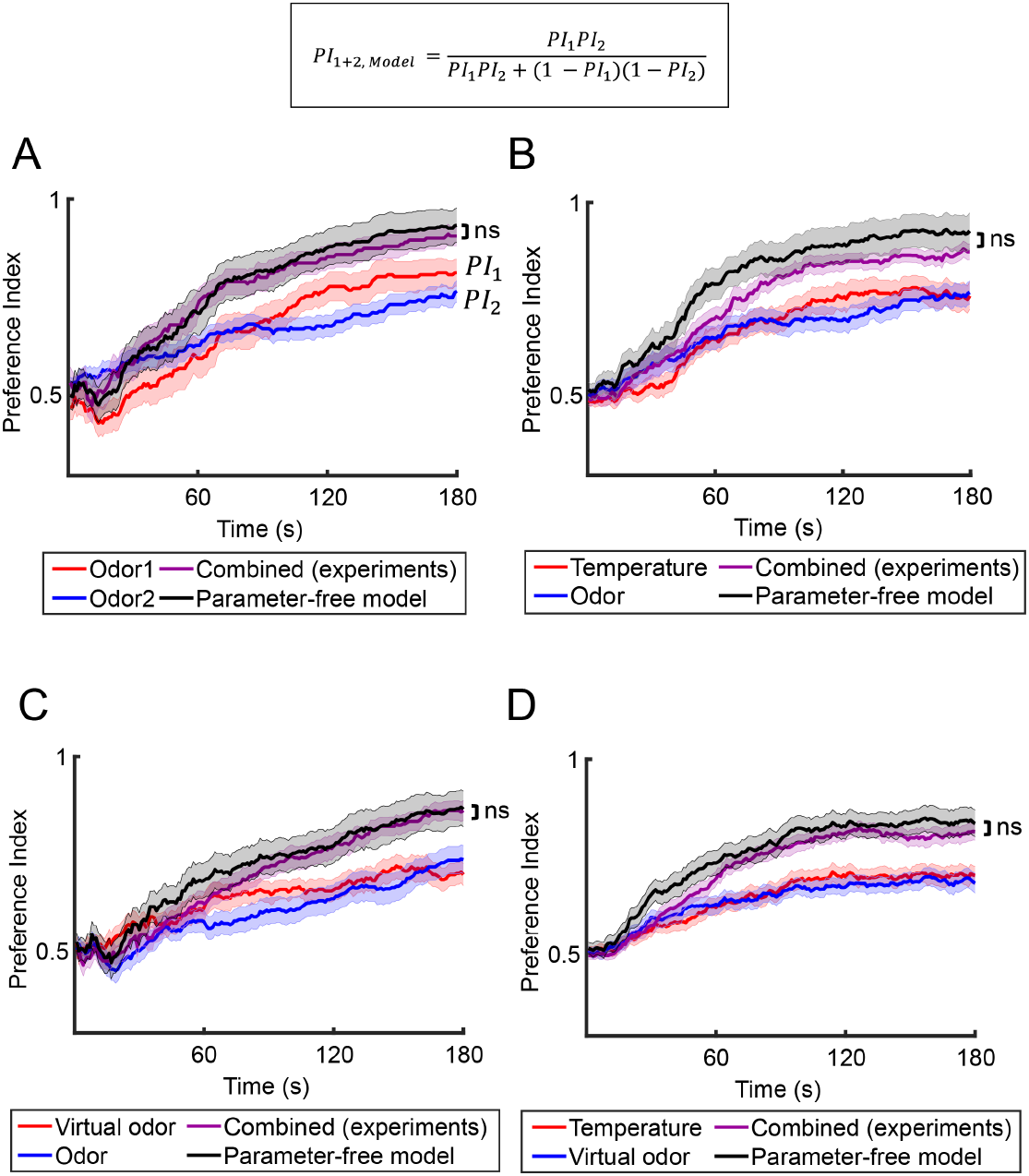
Comparison of the combined preference indices of wild-type larvae with predictions from a parameter-free model for the four configurations outlined in Figure S1 and Figure S2. In all configurations (A-D), there is no significant difference between the final preference indices of the experimental data and the parameter-free model (*t*-test, *p* > 0.05).

**Figure S4.**
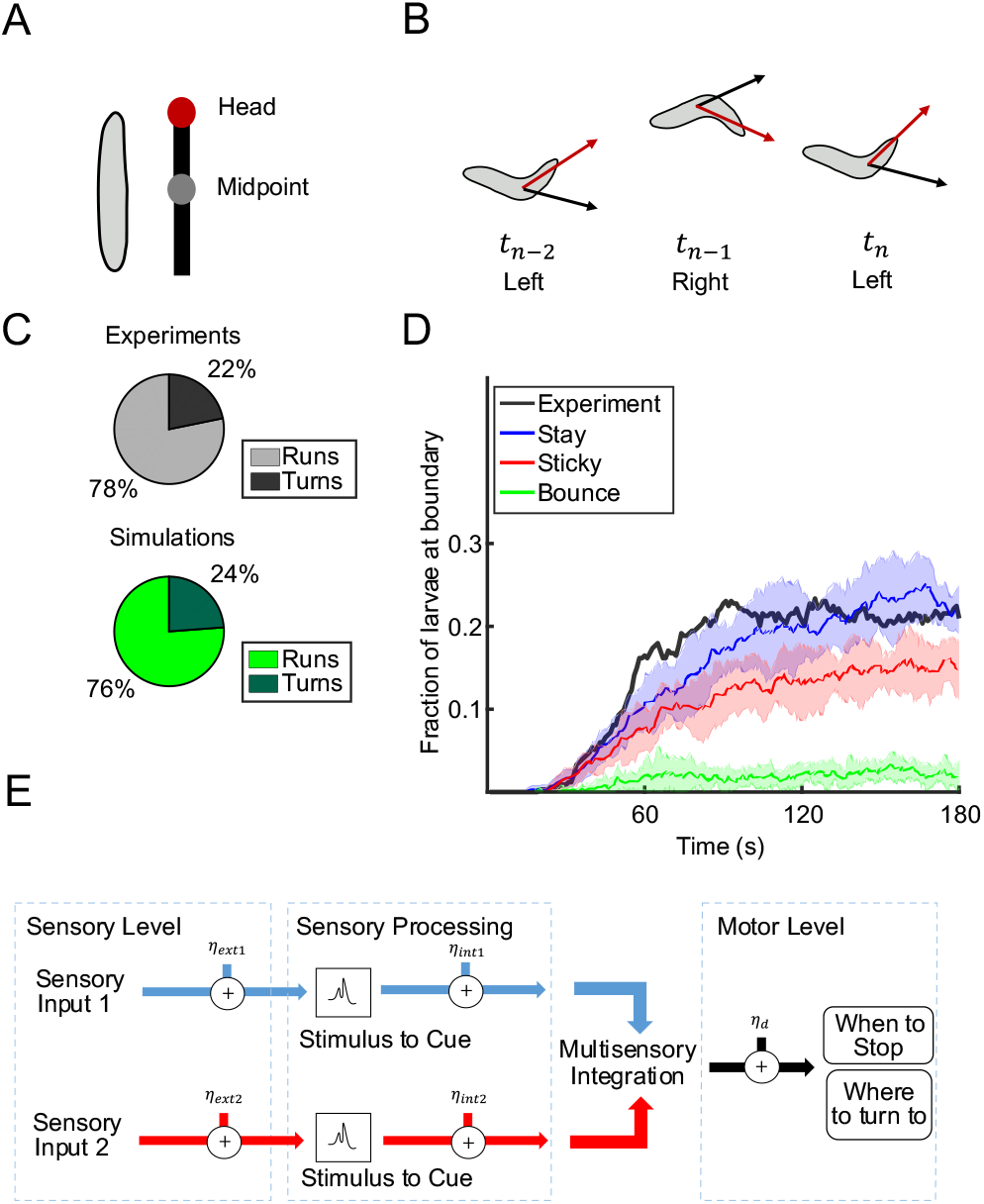
Parameter optimization and performance quantification of the agent-based model for larval navigation. **(A)** Illustration of the framework of the lateral oscillation model (Wystrach et al., 2016) used for the agent based model. The larva is modelled as two segments: the anterior (midpoint to the head) and the posterior (tail). **(B)** The larva alternates between left and right head-casts between every timestep. The black arrow illustrates the direction of motion at the previous timestep while the red arrow is the heading vector at the indicated timestep. **(C)** Ratio of runs and stops observed in real larvae versus in simulations in the absence of stimuli. (n = 100 larvae) **(D)** Simulation results for the fraction of larvae at the walls of the arena for hypothetical boundary conditions tested when designing the agent-based model. Larvae are defined as being at the boundary if they are within one larva-length from the edge of the arena. Lines represent the mean and shaded error bars represent one standard deviation (n = 10 groups of 100 larvae). **(E)** Stages at which noise is added in the agent-based model.

**Figure S5.**
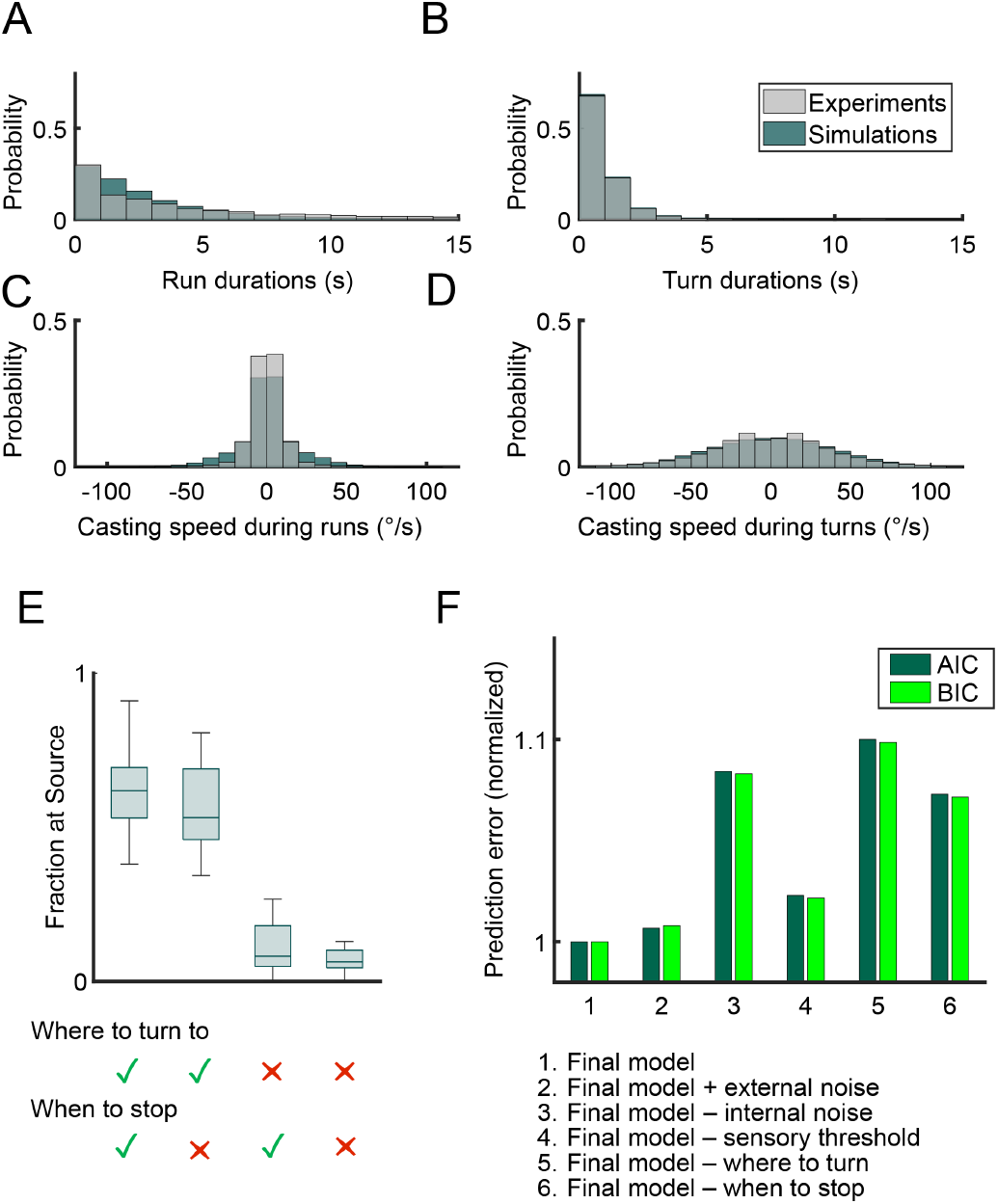
Parameter optimization and performance quantification of the agent-based model for larval navigation. **(A-D)** The histograms compare the behavioral statistics of real larvae to simulated larvae (n = 100 larvae): (A) run durations, (B) turn durations, (C) casting speed during runs, (D) casting speed during turns. (E) Performance of the agent-based model with the removal of its constituent mechanisms (“*where to turn to*”, “*when to stop*”) to direct larvae up gradients. When either mechanism is removed, a smaller fraction of larvae reach the source. (Odor + odor congruent, n = 19 groups of 20 larvae). (F) Justification of model complexity. The plot indicates the change in prediction error as quantified by the AIC/BIC as variables are removed or added to the agent-based model. (Odor + odor congruent, n = 19 groups of 20 larvae)

**Figure S6.**
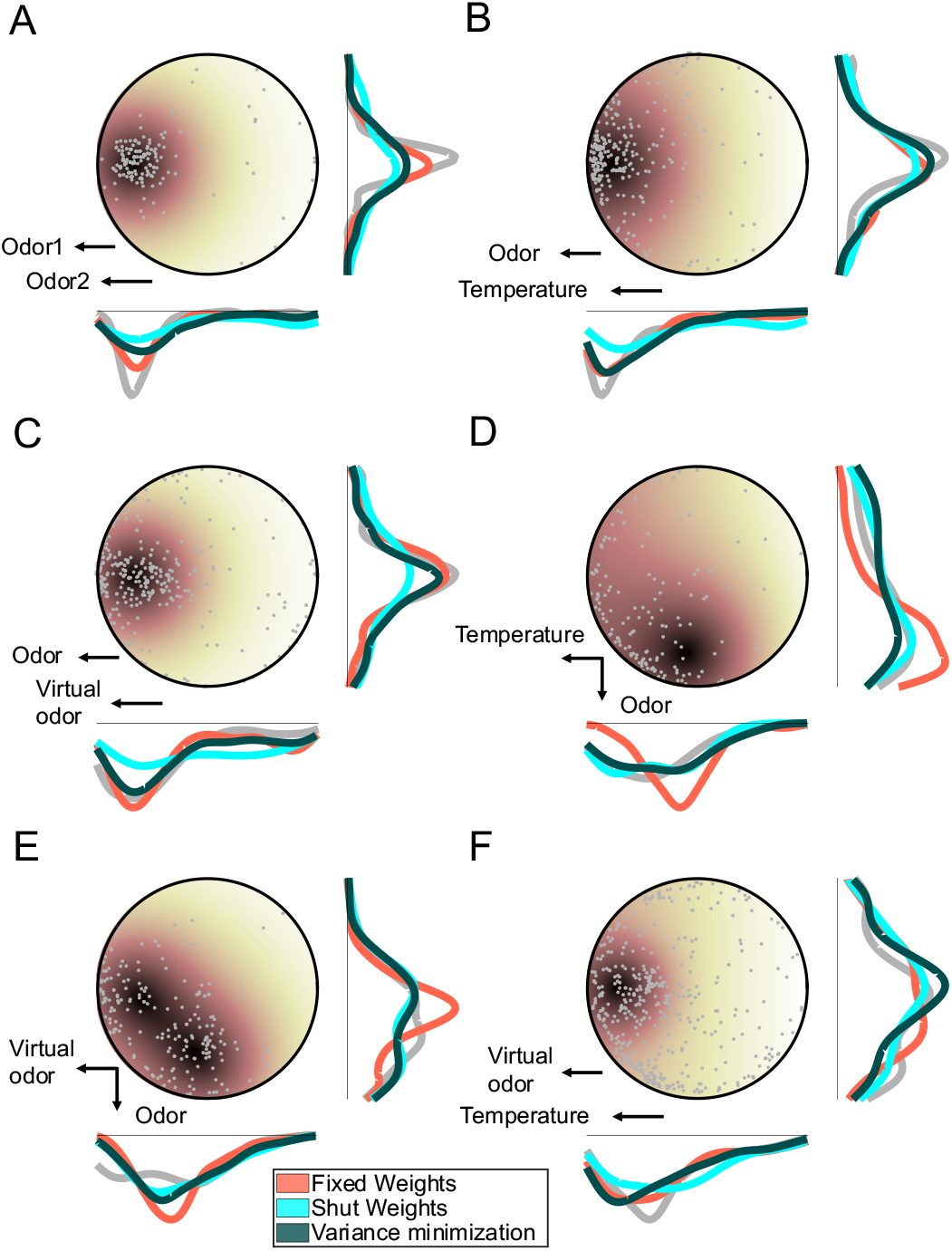
Comparison of final distributions of simulated larvae for each cue-combination rule across different experimental paradigms. **(A)** Odor + odor congruent (odor 1: 1-hexanol, 10^−2^ M; odor 2: ethyl butyrate, 10^−3^ M; n = 19). **(B)** Temperature + odor congruent (odor: ethyl butyrate, 10^−3^ M; temperature: 16-30°C; n = 27). **(C)** Virtual odor + odor congruent (virtual odor: *Or67b*>Chrimson, light 625nm; real odor: ethyl butyrate, 2.5 × 10^−4^ M; n = 30). **(D)** virtual odor + odor conflict (virtual odor: *Or67b*>Chrimson, light 625nm; real odor: ethyl butyrate, 7.5 × 10^−5^ M; n = 20) **(E)** Temperature + odor conflict (temperature: 20-36°C; odor: ethyl butyrate, 2.5 × 10^−4^ M; n = 20). **(F)** Temperature + virtual odor congruent (virtual odor: *Or42a*>Chrimson; temperature: 20-40°C; n = 49).

**Figure S7.**
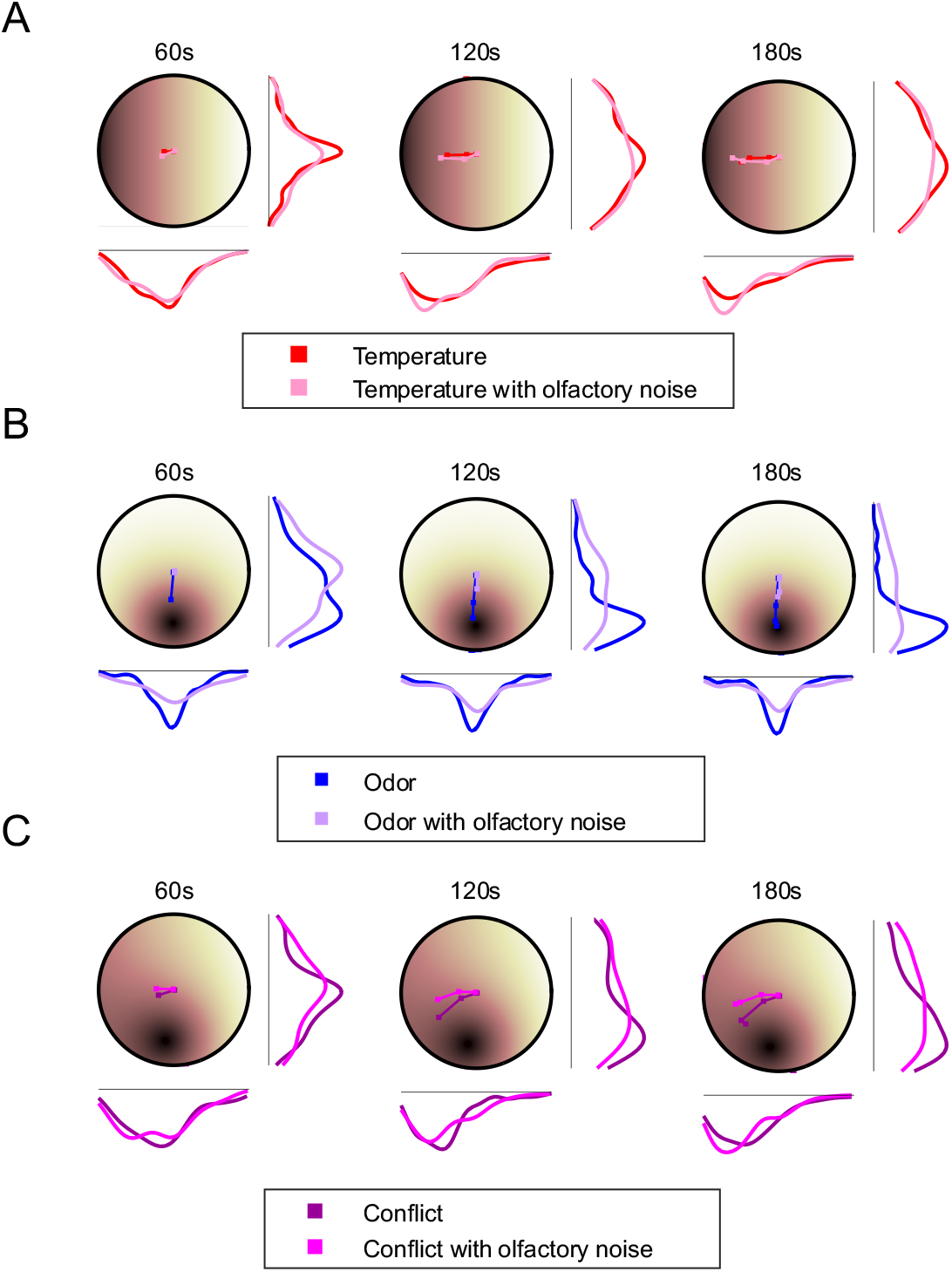
The effect of olfactory noise on navigation in a temperature gradient, an odor gradient, and a conflicting temperature and odor gradient. Each figure shows a comparison of the distributions of real larvae in gradient configurations with and without olfactory noise applied optogenetically (optogenetic olfactory noise: *Or42a*>Chrimson). The mean trajectory of all larvae is shown in the arena over each time interval (60s, 120s, 180s). **(A)** Temperature: 20-36°C, n = 20. **(B)** Odor: ethyl butyrate, 2.5 × 10^−4^ M; n = 20. **(C)** Temperature + odor conflict (Temperature: 20-36°C; odor: ethyl butyrate, 2.5 × 10^−4^ M; n = 20)

**Figure S8.**
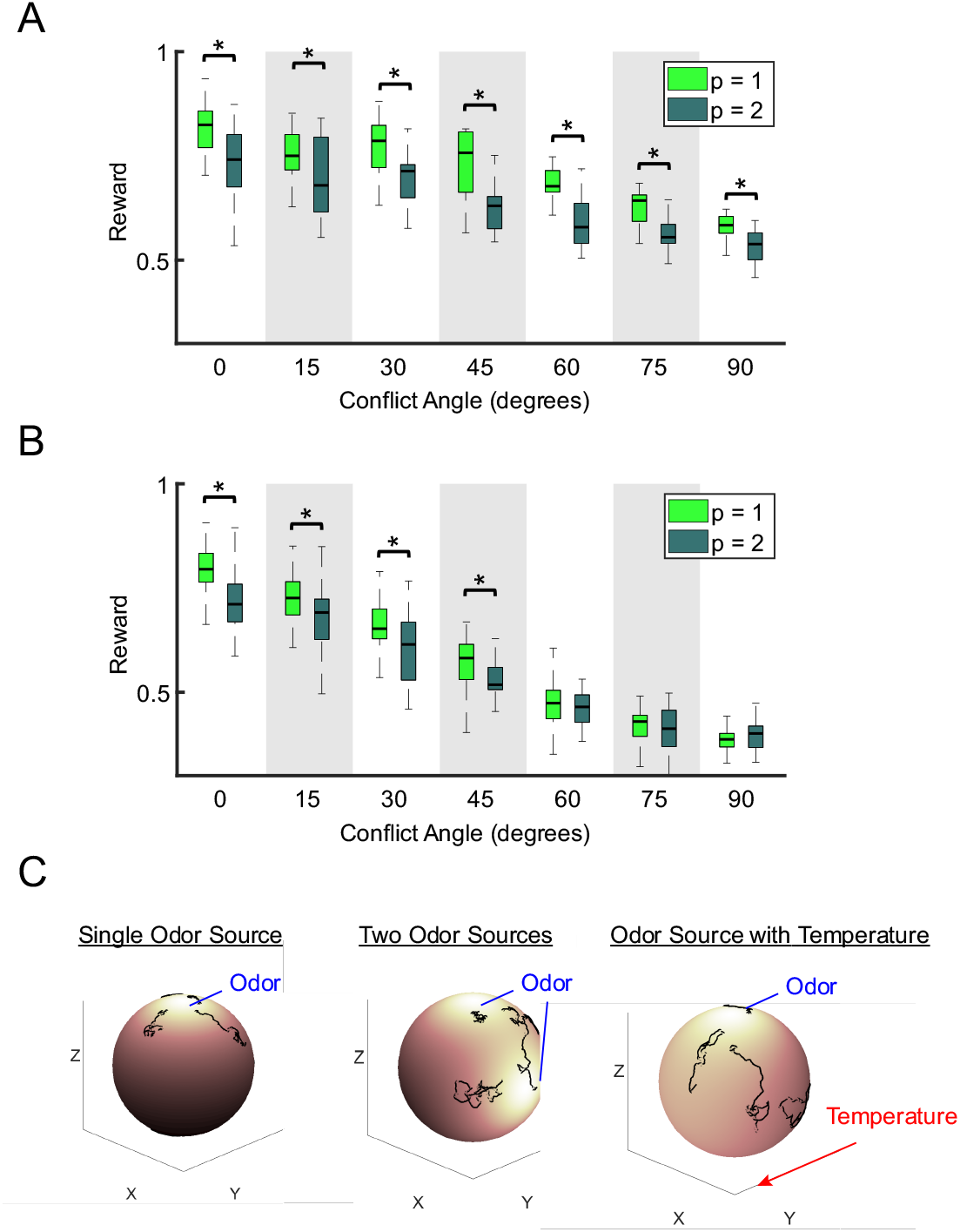
Agent-based model as a testing environment for simulating hypothetical gradient configurations with different conflicting angles (A-B) and 3D configurations (C). **(A)** Virtual odor + odor conflict (n = 19). The mean reward at the end of the simulation is compared between the reward maximization (*p* = 1) and variance minimization (*p* = 2) rules. The asterisk indicates a significant difference by a t-test (p < 0.05) **(B)** Temperature + odor conflict (n = 19). **(C)** Simulations of larvae navigation on the surface of a sphere for different stimulus landscapes (randomly sampled larvae trajectories indicated in black): a single odor source (left), two odor sources (middle), and a single odor source with a linear temperature gradient along the y-axis (right). Color gradient indicates attractiveness of each region (bright = high reward, dark = low reward).

## Supplementary methods

### Parameter-Free Model

*Drosophila* larvae sample the environment to gather information about local odor and temperature gradients through head casts and runs to guide their behavior (Louis et al., 2008). We assume that local information is relatively weak as it is corrupted by fluctuations due to intrinsic noise in the local gradient; thus, the larva needs to accumulate information over time. Two main experimental setups are considered here: one in which two odor gradients are present (one real and another virtual generated by optogenetic stimulation), corresponding to the ‘intramodal’ condition, and another in which an odor and a temperature gradient are present, corresponding to the ‘intermodal’ condition. Mathematically these two conditions can be described with the same formalism, and therefore we do not distinguish them here. We generally use ‘cue 1’ and ‘cue 2’ to refer to either odor or temperature gradients, regardless of the sensory modality used. We will also model the effect of noise injection through optogenetics.

Our model is based on the idea that the larva’s goal is estimating a hidden binary variable *s*, with values −1 and 1, denoting the ‘best location in the world’: if *s* = 1, then the goal location is on the right of the petri dish; if *s* = −1, then the goal location is on the left. The larva estimates this hidden variable by iteratively sampling gradients through the space. We assume that up to time *t*the accumulated evidence for cues 1 and 2 is characterized by sampled gradients Δ*c*_1_ and Δ*c*_2_, respectively. These sampled gradients correspond to the accumulated local sampled gradients, which are lumped together into a single mean-field value. Since sensory observations are noisy due to intrinsic and extrinsic variability, the sampled gradients are corrupted versions of the true

gradients, 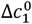 and 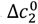 with Gaussian noise. Because both gradients are generated congruently, then we can use the same hidden variable *s* to express 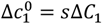 and 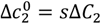, where Δ*C*_*i*_ ≥ 0 are the absolute values of the true gradients 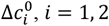, *i* = 1, 2. Therefore *s* represents the sign of the gradient, which points towards the goal location, while 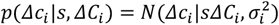 controls the intensity of the gradients. The sampled gradients follow then the equations

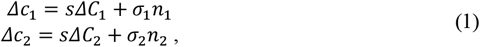

where *n*_*i*_ (*i* = 1, 2) are independent normal random variables with zero-mean and unit variance, and *σ*_*i*_ is the inverse reliability of the *i*-th cue. Control of independence of the fluctuations of the two cues can be achieved in our experiments by using odor and virtual odor gradients.

It is important to emphasize that the primary goal of the larva is to estimate the value of the hidden variable *s* rather than estimating the true values of the gradients Δ*C*_*i*_ through the sampled gradients Δ*c*_1_ and Δ*c*_2_. The variable *s* (the sign of the gradient) specifies the goal location, while the absolute value true gradients are uninformative about the goal location. As the larva estimates the value of the variable *s*, it moves to the estimated goal location. It is important to note that the larva does not have direct access to the true gradient Δ*C*_*i*_ and to the hidden variable *s*. In contrast, in the model the larva has direct access to the inverse reliabilities of each cue through sampling of the noise, as is well documented in other similar scenarios (Ernst & Banks, 2002). This assumption is also supported by our experimental observations.

Errors in the estimated goal location can occur when the two sampled gradients have a different sign with respect to the true location (e.g., when Δ*c*_1_ < 0, Δ*c*_2_ < 0 and *s* = 1). When one of the sampled gradients is positive but the other is negative, then the larva should weigh them according to the reliabilities of each cue. There is a unique way of combining the sampled gradients optimally, the so-called optimal strategy, which we will derive. Our framework is based on Bayesian inference of the hidden variable *s*, which corresponds to the optimal strategy in the sense that the goal location is attained with the highest probability. Given the sampled gradients Δ*c*_1_ and Δ*c*_2_, one can build the posterior probability of the hidden variable *s* and the absolute true gradients as *p*(*s, ΔC*_1_, *ΔC*_2_|*Δc*_1_, *Δc*_2_). Using Bayes’ rule,

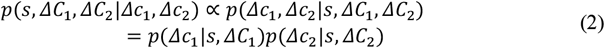

where the proportionality is in relation to *s, ΔC*_1_ and *ΔC*_2_. Since the sampled gradients specify the order of magnitude of the true gradients, and because the true gradients are distributed over several orders of magnitude, we ignore the prior distribution on the true gradients above (effectively, we assume that the prior is flat). In addition, on the right side of the equation we assume that, conditioned on the true gradients and goal location, the fluctuations of the sampled gradients are independent. This is strictly true in our experimental condition in which one gradient is odor and the other is a virtual odor gradient, and they are close-to-independent in other conditions because of the random mixing of odors due to chaotic dynamics in fluids.

Using eq. (1), 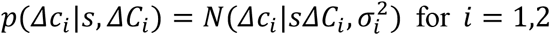 for *i* = 1,2, that is, the density is a Gaussian probability density with mean *sΔC*_*i*_ and variance 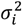. Inserting this expression into eq. (2), we find

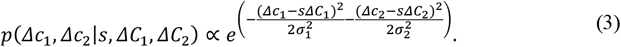

Optimal behavior involves determining the distribution of the hidden variable, but ignoring the absolute values of true concentration gradients, as the latter are not informative about the goal location. Therefore, we are interested in the posterior over the hidden variable *s*, where the absolute values of the gradients are marginalized,

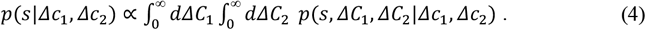

Using eqs. (2-4) and the definition of cumulative Gaussian, 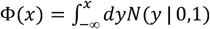, we find

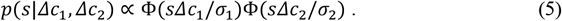

To find a closed expression for *p*(*s*|*Δc*_1_, *Δc*_2_) we approximate the cumulative Gaussians by sigmoid functions, which is known to be an excellent approximation for the best fit parameters (that is, Φ(*x*) is approximated by Φ(*x*) ∼ 1/(1 + *e*^−*αx*^), where *α* is the best fit parameter). Therefore, within this approximation, we can write the probability over *s* as

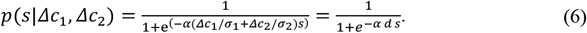

where we have defined the ‘decision variable *d*’

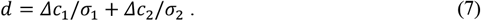

Note that the decision variable weighs the size of the sampled gradients with the reliability of each gradient.

Obtaining the decision variable is one of the central results of this section, as it dictates what the larva should do trial by trial based on the sampled gradients and their reliability. Specifically, when the decision variable is positive, *d* > 0, the probability of *s* = 1 is larger than one half, and therefore optimal behavior dictates moving towards the right. If the decision variable is negative, then optimal behavior dictates moving towards the left. In summary, the decision rule reads:

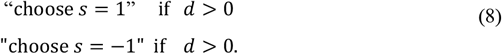

It is important to emphasize that for a larva to follow the optimal behavior it should follow the decision rule in eq. (8). This obviously does not mean that the neuronal circuitry needs to perform explicitly the computation described in eqs. (2-6): all these computations can be bypassed if the decision rule in eq. (8) is hardwired within the neuronal circuits.

The decision rule in eq. (8) is a deterministic rule given the sampled gradients *Δc*_1_ and *Δc*_2_. However, we do not have access to the sampled gradients as measured by the larvae. This means that the value of the decision variable *d* at any particular trial is unknown to us. This implies in turn that we can only know the behavior of the larvae averaged over observations given a predetermined experimental setup, which is characterized by the true gradients 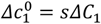 and 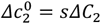. We will take advantage of the fact that, while the true gradients are unknown to the larvae, they are known to the experimenter.

We first note that *d* is the sum of two Gaussian variables, and therefore it is a Gaussian variable. Its mean and variance are respectively

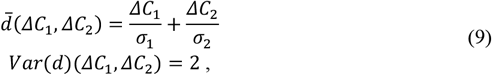

where we assume without loss of generality that the goal location is at *s* = 1. From this expression we can compute the central experimental measurement, the preference index, *PI*. This quantity is defined experimentally as the number of larvae that at time *t* are located on the correct half-side of the petri dish, *s* = 1. We can make a prediction using eq. (9) by noticing that the *PI* is the fraction of times that the decision variable *d* is above zero,

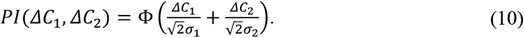

This equation provides a prediction of the preference index when the two gradients are present. Now we can use the same expression to find expressions for the preference indexes for the single-gradient conditions as

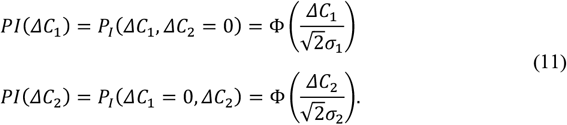

Finally, we can use eqs. (10-11) to obtain the combination rule

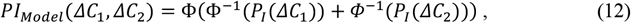

where *Φ*^−1^ (*x*) is the inverse cumulative normal. Thus, using the same sigmoidal approximation of the cumulative Gaussian employed above, we obtain the coarse-grained model given by eq. (2) in the main text. Another important feature of these predictions, which will be exploited later, is that optogenetic stimulation can affect the reliability of each cue in predefined ways. In particular, it should be possible to increase the noise level of cue 1 without affecting the noise level in cue 2. If this happens, then the model predicts that the preference index when only cue 2 is present should remain unchanged in the presence of noise in cue 1. To understand this result, note that in this rule increasing the variance of one signal does not change the total variance of eq. (9), which implies that it is not possible to shut down a cue even if it is very noisy. This is however the optimal thing to do under the above assumption, as the signal is scaled down by the standard deviation of the noise, but gives a different result than the variance weighted combination rule of eq. (7) in the main text. In the main text, *PI*_*Model*_ (*ΔC*_1_, *ΔC*_2_) is denoted as *PI*_1+2,*Model*_.

### Agent-based Model

We model *Drosophila* larvae with an adapted version of an agent-based model developed by *Wystrach et al*. (Wystrach et al., 2016). This model provided a general framework for describing taxis behavior in unimodal stimulus gradients, based on evidence that larvae display continuous lateral oscillations (“head-casts”) of the anterior body during peristalsis. Their work showed that this simple mechanism coupled with the direct sensory modulation of oscillation amplitude could reproduce many taxis signatures observed in larvae. To test different mechanistic hypotheses for cue integration, we build upon this framework to investigate how information can be combined across real odor, virtual odor, and temperature gradients to modulate taxis.

### Lateral Oscillation Model

In our adaptation of the above agent-based model we consider the anterior and posterior body of the larva as two connected segments. The anterior body is modelled as a single segment from the midpoint to the head (Figure S4A). To mimic active sampling, this segment rotates about the midpoint and alternates between left and right rotations between timesteps (Figure S4B), with casting amplitude modulated by the sensory experience. The posterior body on the other hand, is “passive” and assumed to follow the axis of the anterior segment. Larvae are assumed to be uniform in length and move along the anterior heading direction at a constant speed. At any timestep *n* of 1s, this mechanism can be summarized with the following state-update equations:

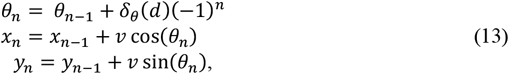

where *θ*_*n*_ is the heading direction of the anterior body relative to the midpoint at timestep *n, v* is the distance travelled in a single time-step, and {*x*_*n*_, *y*_*n*_} is the updated midpoint of the larva. The quantity *δ*_*θ*_ (*d*) is the casting amplitude, which is modulated by a decision variable *d* that is a function of the sensory experience (see below). The constant *v* was estimated based on the average speed observed in larva in the experimental data. In ref. (Wystrach et al., 2016), the amplitudes of the lateral oscillations is modelled as a hard-limit ramp function:

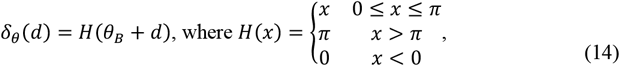

where *θ*_*B*_ is the baseline amplitude of the lateral oscillations in the absence of stimuli (i.e. when *d* = 0). During larval chemotaxis, turning increases during upgradient motion whereas it is reduced during downgradient motion. Accordingly, the decision variable *d* should be negative when moving up a stimulus gradient and positive when moving down a stimulus gradient.

One important feature of our adaptation of the agent-based model that is distinct from Wystrach’s model (Wystrach et al., 2016) is that sensory measurements are sampled at every time-step by a sensor located at the extremity of the larva’s head, which rotates about the midpoint. This allows us to distinguish between head casting during “runs” when the larva is undergoing forward peristalsis and head casting during “stops”, when the midpoint of the larva is stationary. In contrast, the larva in ref. (Wystrach et al., 2016) is modelled as a point agent that rotates on the spot for simplicity. Note that in our model, the position of the larva head is given by:

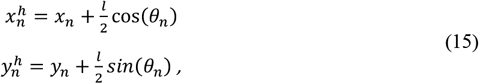

where *l* is the average length of larva at the 3rd instar developmental stage.

### Stopping

For the lateral oscillation model developed in ref. (Wystrach et al., 2016), it was noted that stopping was not essential for chemotaxis except for improving orientation by enabling larger turns in their paths. Thus, this mechanism was ignored as a simplifying assumption and larvae were simulated to run continuously at a fixed speed. However, in order to accurately represent larvae navigation about odor sources in our experimental paradigms, it was necessary to incorporate the mechanism of stopping. We make the following modelling assumptions regarding larvae runs and stops:

1. During runs, larvae move along the anterior heading direction at a constant speed (as before).
2. During stops, larvae remain stationary at the midpoint but are still able to cast the anterior body in either direction.
3. The casting amplitude is larger during stops than during runs.

To capture the behaviors associated with running and stopping in our agent-based model, we assume that larvae not only update their heading direction at each time-step, but also make a decision to run or to stop. Therefore, there are two decisions that must be made at every time-step:

1. *When to Stop*: Should the larva be in a running or stopping state?
2. *Where to Turn*: Given the state of the larva, what adjustment should be made to the current heading?

#### When to stop

We modeled running and stopping in larvae as a binary Markov process, with transition probabilities dependent on the same decision variable *d* (Figure 2E). The transition probabilities between states were given by the following logistic functions:

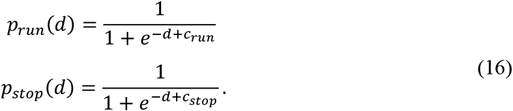

The parameters *c*_*run*_ and *c*_*stop*_ are constants that determine the statistics of running and stopping in the absence of sensory stimuli (i.e. *d* = 0). Using the classification algorithm of the closed loop tracker from ref. (Schulze et al., 2015), we quantified the statistics of running and stopping in unstimulated larvae (Figure S4C). We then used maximum likelihood estimation to fit parameters *c*_*run*_ and *c*_*stop*_ in our model (Figure S5A-D). We verified that the negative binomial distribution of running and stopping durations resulting from the simple Markov model showed a reasonable agreement with actual data.

### Where to turn to

Using experimental data generated with a closed loop tracker (Schulze et al., 2015), we observe differences in both casting amplitude and casting speed in the two states. Given that the dynamics of head casting differ in running and stopping, separate schemes are required to describe the casting amplitude of these two states:

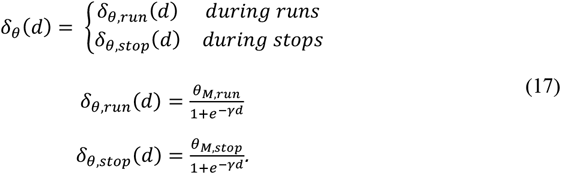

Here, we use a smooth approximation of the hard limit ramp function in ref. (Wystrach et al., 2016). The parameter *θ*_*M*_ can be viewed as a physical constraint on the maximum casting amplitude or head casting speed in running and stopping states. These constants were estimated to fit the physical constraints of the head casting speeds of real larvae. *γ* is a tuning parameter that governs the slope of the ramp and allows for differences in how the decision variable *d* modulates casting amplitude compared to stopping. The resulting head-casting speeds generated by our model were in agreement with real unstimulated larvae from the closed loop tracker.

### Sensory Stimulus

In the present section, we outline the models used to describe the stimulus presented to the larvae. In each experimental paradigm, we presented combinations of dynamic real odor gradients, with static virtual odor gradients and static temperature gradients. At each timestep, we assume that the larva receives a sensory input 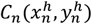 that is dependent on its head position 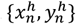 in the assay at timestep *n*.

#### Odor

Two different odors were used in experiments, 1-hexanol and ethyl butyrate. In each experiment, a small odor droplet was placed in an enclosed assay and gradually diffused over the course of three minutes. Since we observed changes in the behavioral response to the odor stimulus over the course of each experiment, we could not assume that the odor gradient was static. Hence, we modeled the evolution of an odor gradient as a diffusion process from a point source as outlined in ref. (Schulze et al., 2015). At timestep *n*, the solution to diffusion partial differential equation is:

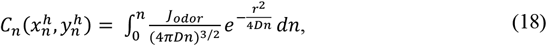

where *r* denotes the Euclidean distance from the larva head to the odor source *r* = 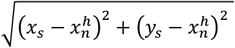 and *J*_*odor*_ is the flux of the odor droplet. *D* is the diffusion coefficient of the odor droplet in air, which differs slightly between *1-hexanol* and *ethyl butyrate*. These values were estimated using the method in ref. (Tucker & Nelken, 1990).

#### T*emperature*

The behavioral experiments feature a linear temperature gradient that varied from *T*_min_ = 16^°^C to a maximum of *T*_max_ = 30^°^C (aversive to larvae). For example, a temperature gradient increasing in the positive x-direction would be given by:

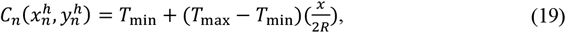

where *R* is the radius of the arena. Under the rearing conditions of the experiments, larvae are drawn to the cooler end of this temperature range.

#### Virtual Odor

In the experiments with real larvae, we passed emitted light from a LED through an exponential filter to create a Gaussian source for optogenetic virtual odor experiments. This is modelled as:

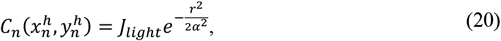

where *r* is again the distance to the source = 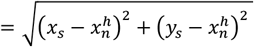, *J*_light_ specifies the intensity of the light stimulus, and *α* is the standard deviation of the Gaussian function. This mathematical fit is supported by measurements of the physical gradient using a photodiode.

#### Sensory Threshold

For experimental conditions involving real odors, we noticed that there was a slight delay in the behavioral response of real larvae at the onset of the experiment. Given that the odor source is introduced in the assay at the same time as larvae, we speculate that the lag in directed behavior is due to the time required for the odor to build up to detectable levels in the arena. To account for this effect, we introduced a sensory threshold parameter *β* such that:

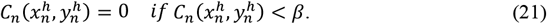

For consistency, we included this threshold as a parameter to be optimized by the framework for all three sensory modalities. However, the effect is significant only for real odors.

### Stimulus to Percept

For each sensory modality presented to the larvae, we assume that the resulting percept (internal intensity representation of the odor) is proportional to relative changes in stimulus strength (Adler & Alon, 2017). Thus, we assume that the perceptual response to the real odor, virtual odor, and temperature gradients will be of the form 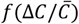, where 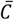 is the background signal level (see eq. (22) below). The validity of this relationship has been established in adult flies (Cao et al., 2016; Kadakia & Emonet, 2019) and it appears to hold for larval olfactory sensory neurons (OSN) that respond to a normalized form of the stimulus derivative (Gomez-Marin & Louis, 2012; Schulze et al., 2015). Although this feature has not been explicitly shown for thermosensation, there is evidence that the behavioral response to an absolute change in temperature increases the larger the deviation from preferred background temperatures (Klein et al., 2015). It was shown further that this process is mediated by cross-inhibition between warming cells and cooling cells (Hernandez-Nunez et al., 2021), activated by positive and negative temperature gradients respectively, and a model was developed to show that the relative contributions of each corresponding signal towards behavior increased as larvae moved away from preferred temperatures. In our experimental paradigm, this would imply that a temperature change of Δ*C* = 1^*°*^C at *T*_max_ = 30^*°*^C would trigger a stronger behavioral effect than an identical change of Δ*C* = 1^*°*^C at the preferred temperature *T*_min_ = 16^*°*^C. We incorporate this perceptually in our agent-based model by rescaling the temperature signal as *C* ← *T*_max_ − *C*. In our simulations, we compute the relative change in stimulus between two consecutive timesteps *n, n* − 1 as the following:

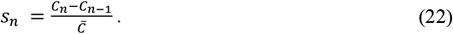

We compute the background signal level as the midpoint between two timesteps. 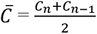. To be able to compare signals from different sensory modalities and stimulus ranges, we define a gain *G* associated with each sensory modality that represents the perceptual sensitivity of larvae. The perceptual (internal) representation of an odor cue, for example, is modelled as:

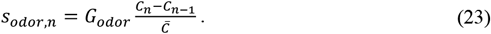

This quantity can be both positive and negative depending on the direction of the sensory gradient. As we do not explicitly model firing rates, we assume that this perceptual representation is encoded by different elements of the peripheral olfactory circuit of the larva. The exact mechanism is unknown; it is not accounted for in the agent-based model.

### Cue Combination

Finally, we model the link between the sensory experience of the larva and its orientation behavior. The mode transitions and casting amplitudes of larva in our agent-based model are described as functions of a decision variable *d*_*n*_, which is dependent on some combination of the sensory modalities perceived by the larva. In subsequent sections, all variables are computed at timestep *n* and we drop the subscript to avoid cluttered notation (e.g. we refer to the decision variable as *d* ≡ *d*_*n*_). We describe the combination of the two different sensory modalities *s*_1_, *s*_2_ using the linear model:

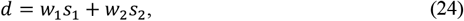

where *w*_1_, *w*_2_ are weights associated with each cue. We hypothesize that larvae may have a bias for one sensory modality over another. Furthermore, we hypothesize that larvae are able to measure the reliability of individual signals when integrating multiple sources of information. We assume that the “reliability” of a sensory signal represented by a time series is inversely proportional to its variance *σ*^2^ (see below). Thus, we test three different plausible weighting strategies:

1. Fixed Weights (FW):

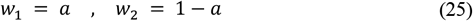
2. Shut Weights (SW):

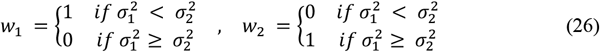
3. Variance Minimization (VM):

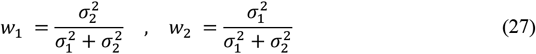

The first weighting strategy proposes that larvae combine cues with fixed preferences that are independent of the signal variance. The latter two strategies imply that larvae are also able to adapt their response according to the estimated variance of the sensory inputs, which has been demonstrated in previous studies (Gepner et al., 2018). The SW strategy assumes that larvae place absolute priority on the cue that is observed to be more reliable. The VM strategy is based on the optimal linear combination rule for minimizing the variance of the combined signal, given certain assumptions (Ernst & Banks, 2002). In the SW and VM models, we assume the larva accumulates sensory evidence over some time window as it navigates the environment and uses this to estimate the variability of each sensory modality. For simplicity, we assume that the variance is estimated through sampling as

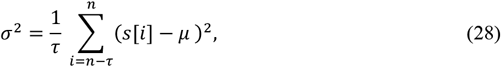

where μ is the sample mean, and *τ* is the time sampling window, which was estimated as *τ* = 11*s* for *Or42a* OSN activation and was shown to be similar in duration for other sensory modalities (Gepner et al., 2018). In the case of a real odor whose concentration is below the detection threshold, the odor would not be perceived as being present and hence the variance *σ* would be assumed to be infinite. This equation assumes that larvae integrate both the temporal variance of the sensory signal itself and self-motion induced spatial fluctuations due to continuous head casting. While it has been suggested that larvae may be able to filter sensory inputs in sync with the frequency of its own peristaltic motion (Gepner et al., 2018), it is unknown how this filtering adapts to motion as the rhythm of head casting is variable and not strictly coupled to peristalsis (Wystrach et al., 2016). Given that it is a weighting of the variances of both channels as ratios that is used to compute cue weights, we assume that the distortions in the estimated variation due to head casting are negligible compared to the true temporal variance of the sensory signal.

### Variance Minimization

For model 3, the decision rule maximizes the reliability of the combined sensory modalities, with the assumption that both gradients originate from a single source (Ernst & Banks, 2002). Let *s*_1_ and *s*_2_ denote the observed cues for attraction from two different gradients, which can be congruent (if the two signs coincide, or incongruent, if the two signs are different). We assume that larvae associate the hedonic value of both gradients in an overall level of attraction, which we denote as *z*. To decide whether to continue in a given direction of motion (heading) or to reorient, larvae infer the latent variable *z* from the observed cues *s*_1_ and *s*_2_. The optimal estimate of the source of attraction *z* can be obtained by applying Bayes rule:

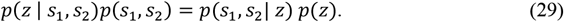

Given that *s*_1_ and *s*_2_ are independent cues as their fluctuations are driven by different physical processes affecting distinct sensory modalities (we neglect joint odor fluctuations due to turbulence, as our assay is far from that regime), we have:

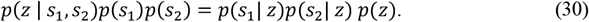

Since *p*(*s*_1_) and *p*(*s*_2_) do not depend on *z*, the variable of interest, we can treat them as proportionality constants:

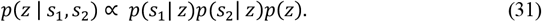

In addition, we assume that the prior *p*(*z*) is flat at every time step, as experiments are performed in an environment that is new to the larvae and there is no evidence that larvae can form spatial memory from previous time steps. We assume that the cues *s*_1_ and *s*_2_ are normal random variables with variances 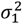 and 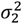. To obtain the optimal estimate of the source of attraction, we calculate the value of *z* that maximizes the posterior probability (maximum a posteriori estimate):

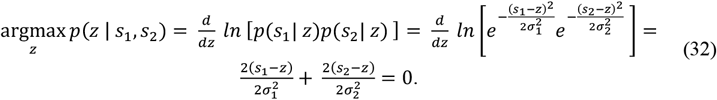

Rearranging, we have:

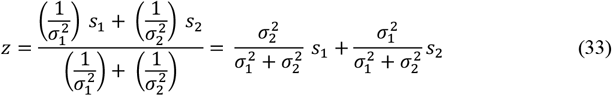

### Reward Maximization

An alternative strategy without assuming a common origin of the two sources is to maximize the expected reward by following each of the two gradients, where reward is defined as the probability that the larva is moving up-gradient. We use the same assumption that the cues *s*_1_ and *s*_2_ are Gaussian random variables with variances 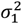 and 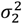. Given any trajectory, the probability that the larva is travelling up-gradient for each of two modalities is 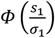 and 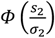, where

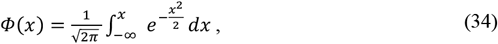

is the standard normal cumulative density function. Assuming that there is an equal preference for reaching either source, the reward of continuing at the current heading is the sum of the probabilities of travelling up-gradient in each of the two sources

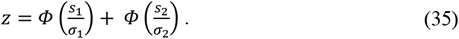

Conversely, the reward of stopping and reorienting is

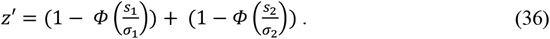

The optimal decision that maximizes reward is therefore to continue at the current heading if *z* > *z*′, and to reorient otherwise. We implement this at the motor level in the agent-based model by defining the decision variable as the reward 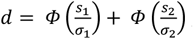, so that the agent will have a low probability of stopping if *d* is large, and will have a high probability of stopping in the opposite case.

### Comparing *p* = 1 (Reward-Maximization) and *p* = 2 (Variance-Minimization) rules

To compare these two strategies, we make several approximations. For maximizing reward, we make the following approximation given *σ*_1_ ≫ *s*_1_ and *σ*_2_ ≫ *s*_2_,

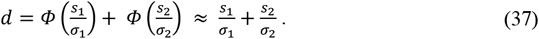

Note that this approximation is identical to eq. (7) in the derivation of the parameter-free model. The *p* = 1 rule corresponds to Reward Maximization. For maximizing reliability, we obtain a different decision variable, namely

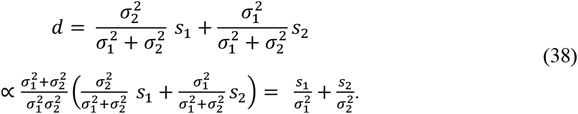

The combination rule with *p* = 2 corresponds to Variance Minimization. In general, we can embed both rules into a single rule with free parameter *p* as

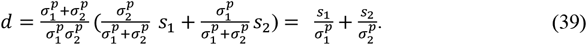

In our simulations, we will optimize the free parameter *p*, as well as compare the *p* = 1 and *p* = 2 rules. We propose to call *p* the *bimodal-contrast parameter*.

### Noise

As we propose that larvae are sensitive to the variance of sensory inputs, an important aspect of this model is to account for noise in the sensory signal. We model noise as Gaussians *η* with zero-mean. For generalizability, we consider noise added at several stages of the flowchart (Figure S4E):

1. Additive external sensory noise: *C*_*n*_+ *η*_ext_
2. Additive internal sensory noise: *s*_*n*_ + *η*_i*n*t_
3. Decision noise: *d* + *η*_d_

The first is additive external noise *η*_ext_ that does not scale with the sensory input. This may be more prominent in experimental paradigms with virtual odor gradients for example, where the noise might result from fluctuations in the action of the LED light on the light-gated ion channel (Chrimson (Klapoetke et al., 2014)). The fixed amplitude light flashes used to perturb the larvae in experimental paradigms with noise can be also modelled with this approach.

The second is additive internal sensory noise *η*_*int*_ due to the assumption that larvae perceive relative changes in stimulus in the agent-based model. Noise that scales with the sensory input would be more plausible for experimental paradigms with real odors, as the fluctuations in odorant molecules tend to fluctuate according to a Poisson distribution, resulting in noise that is dependent on odor concentration.

The third is decision noise, which models the inherent stochasticity of larvae behavior in its mode transitions and variability in casting amplitudes. In our model, we have found similar predictions when incorporating all levels of noise (1 + 2 + 3) and the reduced scheme (2 + 3). While the quality of the predictions may change, we find that the hierarchy of the performance of the weighting strategies does not change with the variations in the framework. This is illustrated in the comparison of AIC and BIC in Figure S5F.

### Optimization Framework

Below is a list of constants used to model larva motion in the simulations:

| Parameter |  | Value |
| --- | --- | --- |
| Run velocity | $v$ | 1.3 mm/s |
| Larva length | $l$ | 3.86 mm |
| Run transition constant | $c_{run}$ | 1.46 |
| Stop transition constant | $c_{stop}$ | 0.16 |
| Maximum casting amplitude during runs | $\theta_{M,run}$ | 0.75 rad/s |
| Maximum casting amplitude during stops | $\theta_{M,stop}$ | 2.93 rad/s |
| Variance sampling time window | $\tau$ | 11 s |
| Decision noise | $\eta_d$ | 0.32 |

These parameters model the movement patterns of foraging 3^rd^ instar larvae in the absence of any stimulus recorded at high spatio-temporal resolution with the closed-loop tracker from ref. (Schulze et al., 2015), and are assumed to be constant across all experimental conditions. The run velocity *v* and larva length *l* were chosen to match the mean observed in wildtype *w*^*1118*^ larva (n = 100 larvae). The parameters *c*_*run*_, *c*_*stop*_, *θ*_*M*, run_, *θ*_*M*, stop_, and *η*_d_ were fit using maximum likelihood estimation as illustrated in Figure S5A-D. The variance sampling time window *τ* was estimated based on the timescale of variance adaptation in (Gepner et al., 2018).

For each experimental paradigm, there are four free parameters associated with each of the two sensory modalities (unimodal conditions):

- *η*_*int*_: Internal additive noise
- *G*: Perceptual gain
- *γ*: Sensitivity to Turning
- *β*: Sensory threshold

Each experimental paradigm has a unimodal condition with each sensory modality presented independently and then a bimodal condition with both sensory modalities presented at the same time. Our approach is to use the data from the unimodal conditions to fit the free parameters of our model, and then use the data from bimodal conditions to evaluate the goodness of fit of the different weighting strategies. Therefore, there are a total of eight free parameters for each experimental paradigm – one set of four parameters for each unimodal condition. We consider the signal and noise of each sensory modality regardless of the test condition (unimodal, bimodal), but we assume that the signal-to-noise ratio is what allows the larva to determine whether a stimulus is present or whether the larva is only perceiving white noise.

To evaluate the goodness of fit of our models, we compared the preference index and the spatial distributions between the experimental data and the simulation.

- **Preference Index:** The preference index (PI) is the fraction of larvae on the preferred side of the arena. The error in the preference index is given by computing the mean squared error between the simulated PI and the experimental PI at different intervals over the course of the experiment.
- **Spatial Distribution:** We use the Kullback-Leibler (KL) divergence to compare the error between the simulated and experimental spatial distributions over the entire course of the experiment. The X and Y dimensions are considered separately when computing the KL divergence.

Because the preference index only measures the fraction of larvae that are on the preferred side of the arena, we find that the spatial distributions give a more accurate representation of the quality of fit. All parameter fitting was performed using the Global Optimization Toolbox in MATLAB. Below is a list of the median parameter values for each experiment across different tested bimodal contrast coefficients *p*:

| Experiments |  |  | Parameters |  |  |  |  |  |  |
| --- | --- | --- | --- | --- | --- | --- | --- | --- | --- |
| Gradient 1 | Gradient 2 | Orientation | $G_1$ | $\eta_{int,1}$ | $\beta_1$ | $G_2$ | $\eta_{int,2}$ | $\beta_2$ | $\gamma$ |
| **Odor**<br>**(1-hexanol)** | Odor<br>(ethyl butyrate) | Congruent | 4.90 | 0.13 | 0.28 | 5.75 | 0.18 | 0.68 | 1.09 |
| **Virtual odor** | Odor<br>(ethyl butyrate) | Congruent | 3.78 | 0.47 | 5.54 | 4.90 | 0.26 | 0.56 | 1.05 |
| **Temperature** | Odor<br>(ethyl butyrate) | Congruent | 7.06 | 0.37 | 2.23 | 6.70 | 0.24 | 0.27 | 0.98 |
| **Temperature** | Odor + noise<br>(ethyl butyrate) | Congruent | 7.06 | 0.37 | 2.23 | 6.70 | 0.40 | 0.27 | 0.98 |
| **Temperature** | Virtual odor | Congruent | 2.00 | 0.55 | 5.10 | 2.75 | 0.57 | 5.13 | 1.20 |
| **Virtual odor** | Odor<br>(ethyl butyrate) | Conflict | 4.01 | 0.33 | 5.03 | 6.33 | 0.23 | 0.29 | 0.97 |
| **Temperature** | Odor<br>(ethyl butyrate) | Conflict | 5.46 | 0.26 | 2.88 | 6.62 | 0.15 | 0.18 | 0.96 |
| **Temperature** | Odor + noise<br>(ethyl butyrate) | Conflict | 5.46 | 0.26 | 2.88 | 6.62 | 0.45 | 0.18 | 0.96 |

### Simulating Wall (Boundary) Conditions

Since the arena is small, one last component of our model is accounting for larvae behavior at the edges for the arena. We noted that a significant fraction of larvae remained close to the arena boundary (its wall), particularly in conditions with a linear temperature gradient. We considered several possibilities if a larva’s path is obstructed by the arena wall (Figure S4D):

1. The larva remains stationary in a stopping state as long as its position at the next timestep is outside the bounds of the arena.
2. The larva moves tangent to the edge of the arena at a velocity *v*_*edge*_ = cos(*ψ*) *v*, where *v* is the larva’s original speed, and *ψ* is the angle between the larva’s heading direction and the direction tangent to the arena.
3. The larva “bounces” off the edge of the arena at the angle of incidence (ballistic collision model).

Through numerical simulations, we found that the first approach is the closest representation of the behavior observed in our experimental data based on the stopping statistics of larvae at the boundary.

### *Fraction-at-Source* and *Reward* Metrics

The *“Fraction at Source”* is defined as the number of larvae within bounded regions near the peak of the gradients divided by the total number of larvae:

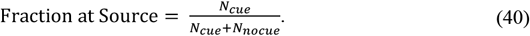

For odor configurations, this bounded region is defined as an area within radius *r* of the source. For temperature configurations, the bounded region associated with the comfortable (targeted) temperature is any location *x* < *r*, where *x* = 0 corresponds to the leftmost, coolest side of the arena. The radius *r* was chosen such that the areas of the bounded regions were identical for both odor and temperature configurations (*r* = 1.8*cm*). The *“reward”* for each sensory modality is defined as the mean perceived sensory experience of all larvae relative to the peak sensory experience in the arena. In the bimodal condition, the reward is calculated as the average reward across both sensory modalities. For *N*_*j*_ number of sensory modalities, the reward is given by:

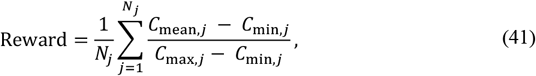

where *C*_mean,*j*_ is the mean sensory experience of all larva for sensory modality *j*, while *C*_min,*j*_ and *C*_max,*j*_ denote the least and most preferred sensory experience in the arena respectively for sensory modality *j*.

### Model Selection with AIC/BIC

The prediction error for the AIC/BIC (Akaike, 1998; Schwarz, 1978) was computed for the Variance Minimization rule across all bimodal experimental paradigms:

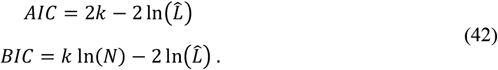

Where *k* is the number of model parameters, *N* is the number of simulated larvae for each experimental paradigm, and 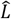 is the likelihood function given the actual observed spatial distributions of larvae. In each model variant, one component of the model was added/removed,

and the model parameters were re-optimized. The resulting prediction error was then compared to that of the final model. All variations of the model resulted in a higher prediction error, as shown in Figure S5F.

